# Probabilistic index models for testing differential expression in single cell RNA sequencing data

**DOI:** 10.1101/718668

**Authors:** Alemu Takele Assefa, Jo Vandesompele, Olivier Thas

## Abstract

Single-cell RNA sequencing (scRNA-seq) technologies profile gene expression patterns in individual cells. It is often of interest to test for differential expression (DE) between conditions, e.g. treatment vs control or between cell types. Simulation studies have shown that non-parametric tests, such as the Wilcoxon-rank sum test, can robustly detect significant DE, with better performance than many parametric tools specifically developed for scRNA-seq data analysis. However, these rank tests cannot be used for complex experimental designs involving multiple groups, multiple factors and confounding variables. Further, rank based tests do not provide an interpretable measure of the effect size. We propose a semi-parametric approach based on probabilistic index models (PIM) that form a flexible class of models that generalize classical rank tests. Our method does not rely on strong distributional assumptions and it allows accounting for confounding factors. Moreover, it allows for the estimation of the effect size in terms of a probabilistic index. Real data analysis demonstrate that PIM is capable of identifying biologically meaningful DE. Our simulation studies also show that DE tests succeed well in controlling the false discovery rate at its nominal level, while maintaining good sensitivity as compared to competing methods.

## 1 INTRODUCTION

Single cell RNA sequencing (scRNA-seq) technologies have become powerful tools to study the dynamics of gene expression at unprecedented high resolution [1, 2]. The data produced by such scRNA-seq technologies represent measures of transcript/gene abundance in individual cells. In the field of biology and medicine, it is actively used in various applications including delineating population diversity, tracing cell lineages, classifying cell types, and genomic profiling of rare cells [2]. For example, analysis of scRNA-seq data is of paramount interest in studying diseases, such as cancer, to characterize intra-tumoral heterogeneity and to classify cancer cell sub-populations [2]. Of note, scRNA-seq data are known to be associated with particular issues that challenge ordinary data analysis pipelines [3, 4]. Such issues include high abundance of zero counts, over-dispersion, and complex expression distributions (multimodal distributions) [5, 6, 7]. These issues are the result of either intrinsic biological variation or technical artifacts involved in various steps of the data processing pipeline [1, 6, 8]. Yet, the high level of noise can to some extent be compensated by the large number of cells processed in a scRNA-seq study [7].

One of the most common gene level analyses is testing for differential expression (DE). DE analysis aims at identifying a set of genes with different distributions of expression across populations of cells. Yet, the high magnitude of noise in scRNA-seq data challenge DE analysis in terms of controlling the type-I error rate, and hence the false discovery rate [9]. Various statistical methods for testing DE have been introduced. Many of them start from the assumption that the distribution of the data can be described by a particular parametric probability distribution. Among these, (zero inflated) negative binomial [10, 11] and two-component mixture distribution are common [12]. Parametric methods provide a flexible and straightforward modelling framework focusing on the difference in the mean expression across groups. Nevertheless, it is well known that the performance of parametric methods depends on the validity of the distributional assumptions, as well as on the precision of the parameter estimators. Simulation studies have demonstrated that non-parametric tests, such as the Wilcoxon rank sum test and SAMSeq [13], are equally useful for robustly testing DE with competitive performance in terms of sensitivity and false discovery rate (FDR) control [9, 14, 15]. Unfortunately, the rank tests available for scRNA-seq have a limited scope. First, they cannot be used for experiments involving comparison of multiple populations of cells or more than one factor of interest [9]. For example, Xin et al. [16] demonstrated cell type-specific DE between type-2 diabetics and controls while adjusting for subject specific characteristics such as gender and ethnicity. This design involves two main factors (disease status and cell type), their interaction effect and two additional nuisance factors, in which case simple rank-based tests cannot be used. Second, classical rank tests focus on hypothesis testing and they do not provide an estimate of an interpretable effect size measure. In this paper, we propose a new and flexible method that broadens the scope of rank-based tools and alleviates their limitation while retaining their robustness for testing DE in scRNA-seq data.

We propose a semi-parametric method based on Probabilistic Index Models (PIM) [17, 18], for testing DE in scRNA-seq data. PIMs entail a large class of semi-parametric models that can generate many of the traditional rank tests, such as the Wilcoxon-Mann-Whitney test and the Kruskal-Wallis test [17, 18]. These models can be seen as the rank-equivalent of the generalized linear models (GLM). PIM methods do not rely on strong distributional assumptions and they inherit the robustness of rank methods. Moreover, PIMs come with parameters that possess an informative interpretation as effect sizes. PIMs can be used for complex experimental designs involving multiple groups of cells, and multiple discrete or continuous factors. PIMs form a regression framework and enable accounting for confounding factors that drive unwanted variations such as library size difference, batch, and cell cycle stage. Consequently, PIMs can be applied on the raw count data without the need for a normalization preprocessing step. Of note, it is also possible to integrate PIMs with other data preprocessing methods, such as normalization, imputation, and removal of unwanted variation.

## 2 METHODS

Let (*Y*_*i*_, *X*_*i*_), *i* = 1, 2, …, *n*, be a set of *n* sample observations with *Y* the outcome variable and *X* the corresponding *p*-dimensional covariate. In a GLM, the conditional mean E(*Y |X*) is modelled as a function of covariates through an appropriate link function. In contrast, a PIM models the conditional probability

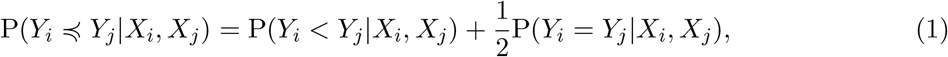

where *Y*_*i*_ | *X*_*i*_ and *Y*_*j*_ | *X*_*j*_ have conditional distribution functions *F* (*⋅*; *X*_*i*_) and *F* (*⋅*; *X*_*j*_), respectively, with *F* further unspecified. The probability in (1) is known as the *probabilistic index* (PI) [17]. For a link function *g*(.), PIM is defined as

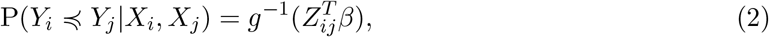

where *β* is the regression parameter and *Z*_*ij*_ is a function of the pair of covariates *X*_*i*_ and *X*_*j*_. The latter is very often *Z*_*ij*_ = *X*_*j*_ − *X*_*i*_. Similar to a GLM, the systematic component is restricted to 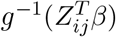 with an appropriate link function *g*(.). Throughout this paper, we use the *logit* link function (i.e. 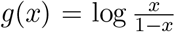). To obtain the parameter estimates, say 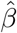, the estimation method uses *pseudo observations I*_*ij*_ = *I*(*Y*_*i*_ < *Y*_*j*_) + 0.5I(*Y*_*i*_ = *Y*_*j*_), where I(.) is the 0/1 indicator function. The pseudo-observations *I*_*ij*_ take values in {0, 0.5, 1}, and, like rank-based methods, they depend only on the ordering of the outcomes. It is also shown that the estimator 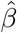 is asymptotically normally distributed and a consistent estimator of its variance is available. We refer readers to [17], [18] and the supplementary file for further details about PIMs.

### 2.1 PIMs for testing differential gene expression

We propose PIMs for testing DE across experimental/biological conditions in scRNA-seq data. To illustrate how PIMs can be implemented for testing DE in scRNA-seq data, we first focus on a simple scenario with two groups of cells. Let *Y*_*gi*_ denote the normalized gene expression of gene *g* = 1, 2, …, *G* in cell *i* = 1, 2, …, *n*. Later, we extend the model so that it can work directly on the raw counts. Let covariate *X* = *A* be the group indicator, such that *A*_*i*_ = 1 if cell *i* is in one group and *A*_*i*_ = 0 if the cell is in the other group. We specify a PIM using the logit link function as

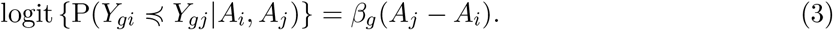

where the indices *i* and *j* refer cell *i* and *j* with associated grouping factor *A*_*i*_ and *A*_*j*_, respectively. The parameter *β*_*g*_ ∈ ℝ represents the effect of the factor *A* on the PI of the outcome. This effect is expressed as *P* (*Y*_*gi*_ ≼ *Y*_*gj*_|*A*_*i*_ = 0, *A*_*j*_ = 1) = exp(*β*_*g*_)/(1 + exp(*β*_*g*_)). Once its estimate 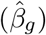 is obtained, the PI can be estimated as

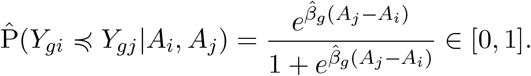

If there is a strong evidence that gene *g* is DE, then the estimated PI 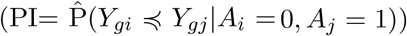 becomes close to 1 (if the gene expression is higher in the *A* = 1 group) or 0 (if higher in the *A* = 0 group). Under the null hypothesis (no DE), the estimated PI is expected to be 0.5, indicating that there is a 50% chance that the expression of gene *g* in a randomly selected cell from the *A* = 0 group is lower than that of a randomly selected cell from the *A* = 1 group (and vice versa). In other words, the PI indicates to what extent the distributions of the gene expression in the two groups are well distinguished. Thus, in this simple setting, the PI is equivalent to the area under the receiver-operating-characteristics curve (AUC) in a classification problem. This is graphically illustrated in Figure 1.

**Figure 1:**
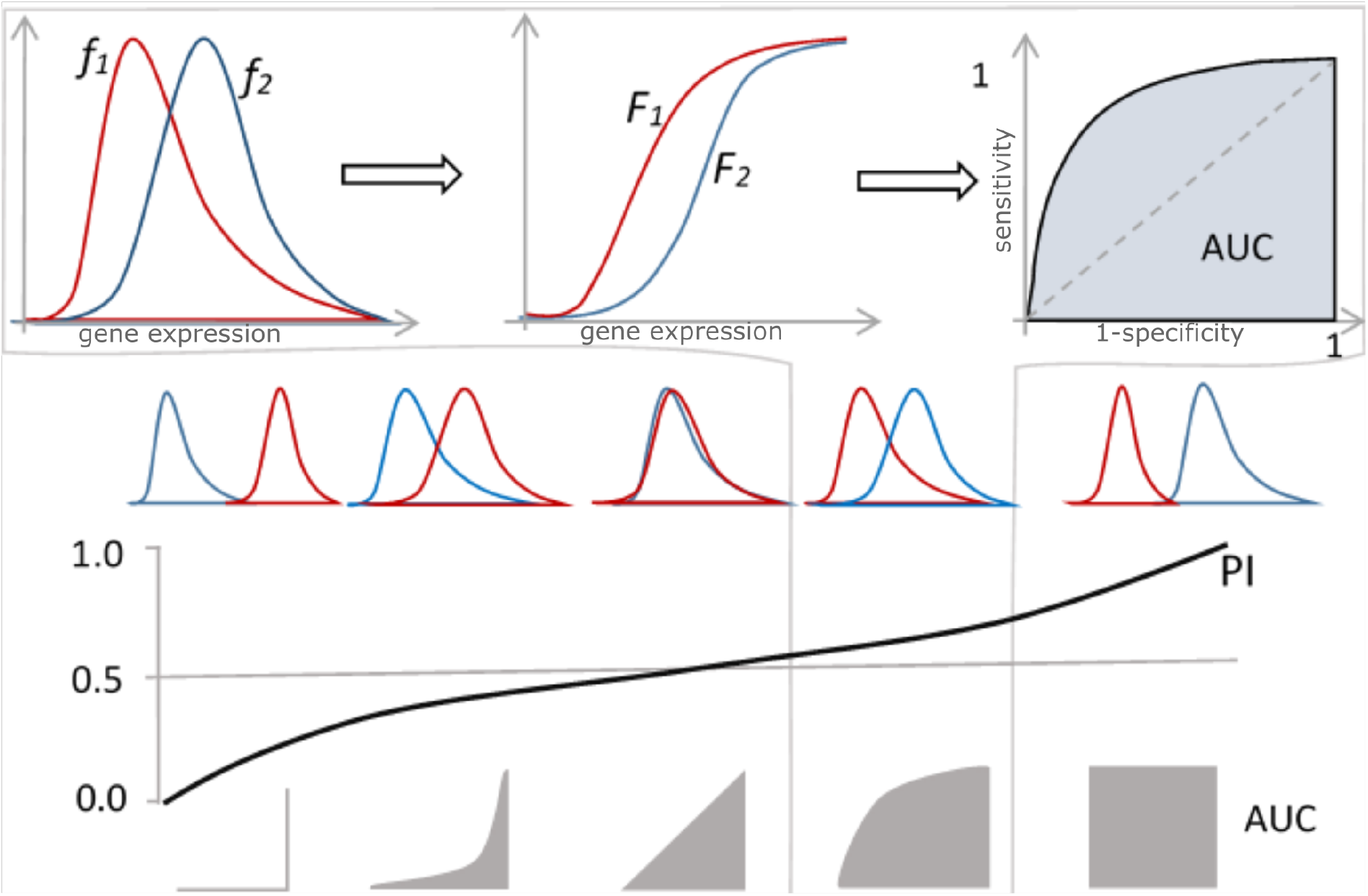
Interpretation of the Probabilistic Index (PI). The PI indicates to what extent the gene expression level distributions of two groups are different. In other words, the PI is equivalent to the area under the receiver-operating-characteristics curve (AUC) in a classification problem. Let *Y*_*i*1_ and *Y*_*i*2_ be the gene expression levels in group 1 and 2 with *F*_1_ and *F*_2_ their corresponding distribution functions, and *f*_1_ and *f*_2_ their density functions. Assume we want to classify cells into two groups based on the expression level of a given gene (high or low expression at a particular threshold t). Thus, the PI =P(*Y*_*gi*_ ≼ *Y*_*gj*_) is equivalent to AUC for classifying a cell into group 2 (if expression> *t*) and group 1 (if expression< *t*) for all possible values of *t*. If the gene expression level is higher in group 2 than in group 1, then PI and AUC will be close to 1. Conversely, if the gene expression is higher in group 1 than in group 2, then PI and AUC will be close to 0. If there is no difference in expression, then both PI and AUC will be approximately 0.5

Statistical hypothesis testing can be performed with asymptotic Wald-type test of Thas et al. [17]. For example, for the model in (3), the null hypothesis of no DE can be formulated as *H*_0_: *β*_*g*_ = 0. This is equivalent to testing P(*Y*_*gi*_ ≼ *Y*_*gj*_|*A*_*i*_ = 0, *A*_*j*_ = 1) = 0.5. Applying this test to every candidate gene results in a vector of raw *p*-values, to which the usual FDR control procedures, such as the Benjamin and Hochberg [19] approach, can be applied. In the subsequent sections we will demonstrate that the *p*-values satisfy the assumptions required by such procedures.

### 2.2 Normalization and removal of unwanted variation

In the previous section, we assumed that the gene expression data were normalized per cell, using e.g. simple methods, such as transcripts per kilobase million (TPM) or counts per millions of reads (CPM) [20], or other (bulk RNA-seq) normalization methods that involve the calculation of a global scaling factor to normalize gene expression levels in each sample by a single constant. Importantly, typical scRNA-seq data are often confounded by several sources of unwanted variation among cells, such as library size, batch, cell size, cell cycle, and cell quality level that make cells not directly comparable [4]. Consequently, normalization of scRNA-seq data at best requires accounting for these nuisance factors, and the meaning of this scaling factor is broader than in bulk RNA-seq data in which normalization is generally needed to remove library size differences [4, 21]. Moreover, the effect of such factors (e.g. cell cycle and batch) can be gene dependent [6]. Cole et al. [4] and Risso et al. [22] demonstrated that regression-based normalization is effective in addressing the technical factors in scRNA-seq data. PIMs allow for the incorporation of such known sources of variation. In this way, we can test for DE, while controlling for various technical factors at a gene level. In particular, let **W** be an *n × q* matrix of *q* cell level technical factors, then the PIM in (3) can be expressed as

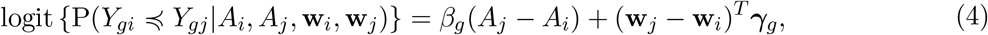

where **w**_*i*_ and **w**_*j*_ are *q*-dimensional vectors of technical factors for cells *i* and *j*, respectively, and ***γ***_*g*_ is a *q*-dimensional parameter vector. In this model, *Y*_*gi*_ is the observed expression of gene *g* in cell *i*. In particular, the adjusted effect of the group factor *A* on the PI becomes

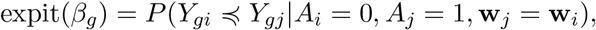

where expit(*x*) = exp(*x*)*/*(1 + exp(*x*)). Thus, testing for DE between *A* = 0 and *A* = 1 is adjusted for the effect of the known nuisance factors **W**.

PIMs are flexible to be used for testing DE in a scenario with a multi-class factor or multiple discrete or continuous factors. For multiple factor studies, PIMs also allow for testing for interaction effects in which case one can test for the global effect or selected contrasts.

### 2.3 Cox proportional hazard model approximation of PIM

The standard parameter estimation methods for PIMs involve comparison of all outcome pairs, increasing the computation time quadratically with the sample size. To improve scalability and computational efficiency for datasets with large number of cells (for example, *n >* 500), we propose the Cox-proportional hazard (Cox-PH) model [23] translation of PIMs. The Cox-PH model implies a PIM with logit link function and the covariates specification as **Z**_*ij*_ = **X**_*j*_ − **X**_*i*_ [17, 18, 24] (see Supplementary File for the details). While the Cox-PH model comes at the cost of the assumption of proportional hazard (PH), PIMs are less restrictive than the Cox-PH models, and results from PIM are still valid when the PH assumption does not hold [24]. Our empirical assessment of the PH assumption in scRNA-seq data for the model in (4) shows that the PH assumption holds for at least 91% of the genes in dataset B, and for 82.5% of genes in dataset A (Supplementary Table S1, see the next section for the description of the datasets).

### 2.4 Single cell RNA-seq dataset

To benchmark PIM and demonstrate its application, we used two scRNA-seq datasets, from two different experimental protocols (see Table 1).

**Table 1:** Summary of the scRNA-seq datasets

| data | protocol | gene<br>expression<br>measure type | number<br>of cells | number<br>of genes | library size*<br>(min-max)<br>in $10^5$ |
| --- | --- | --- | --- | --- | --- |
| A | SMARTer/C1 | read counts | 83 | 22,078 | 3.21 - 12.36 |
| B | Chromium | UMI** | 7149 | 18,639 | 0.04 - 0.21 |

\* library size per cell
\*\* UMI=unique molecular identifier

The first dataset (A) is a cellular perturbation experiment on the C1 instrument [25] (GEO accession GSE119984). This total RNA seq dataset contains 83 NGP neuroblastoma cells of which 31 were treated with nutlin-3 and the other 52 cells were treated with vehicle (controls). The second scRNA-seq dataset (B) contains the same neuroblastoma cells under nutlin-3 treatment and control, except that the data are generated using the Chromium (10X Genomics) instrument with 3’ end sequencing (GEO accession GSE144931). We applied PIM and other competitor tools for testing DE between the treatment and control groups of cells in these two datasets.

### 2.5 Simulation methods

We implemented three simulation methods, each starting from other underlying assumptions. In the simulation study, the PIM specification in (4) with *w*_*i*_ the log-library size of cell *i* was compared with the competitor tools.

#### Mock comparison

This simulation involves random assignment of cells from a single condition/population to two mock groups. Since we do not expect DE between mock groups, it enables the evaluation of controlling the type-I error rate (also known as false positive rate). In particular, cells within the control group (vehicle) in the two scRNA-seq datasets are randomly split into two mock groups between which no DE is expected. Any rejection of the null hypothesis at the 5% nominal per-comparison error rate (PCER) level is considered as false positive. From running 100 independent simulations, the actual type-I error rate for each gene is approximated as the fraction of tests with un-adjusted *p*-value less than 5%.

#### Splat simulation – gamma-Poisson hierarchical model

This simulation algorithm makes use of a gamma-Poisson hierarchical model [26]. In particular, the mean expression level of gene *g* in cell *i*, *λ*_*gi*_, is sampled from a gamma distribution (i.e. *λ*_*gi*_ ∼ Γ (*α*_*gi*_, *β*_*gi*_)). Subsequently, the read counts *Y*_*gi*_ for gene *g* in cell *i* are sampled from a Poisson distribution (i.e. *Y*_*gi*_|*λ*_*gi*_ ∼ Poisson(*λ*_*gi*_). The hyper-parameters *α*_*gi*_ and *β*_*gi*_ are chosen accounting for the desired library size in cell *i* and the mean-variance trend across genes. Afterwards, Splat uses the logistic function for the observed relationship between the mean expression of a gene and the proportion of zero counts to add excess zeros representing technical noise (also called dropouts). It is implemented using the *splatter* R Bioconductor package (version 1.6.1)[26]. The gamma-Poisson distribution is a particular parametrization of the negative binomial (NB) model, and hence the Splat simulation is equivalent to simulation from a NB model. Splat uses model parameters estimated from a real single cell RNA-seq data set. To add a set of DE genes, the mean expression of randomly selected genes are multiplied by a factor, known as the fold-change.

#### SPsimSeq simulation

Since the (zero inflated) NB distributional assumption for scRNA-seq data is debatable [11, 27], the results of the parametric simulations should be interpreted with care. SPsimSeq [15] is a semi-parametric simulation method for bulk and scRNA-seq data. It makes use of no specific parametric distribution for the gene expression levels. Instead, it generates data from a more realistic distribution, which is estimated from a real source dataset by the semi-parametric method of [28] and [29]. In a first step, the log-CPM values from a given real data set are used for semi-parametrically estimating gene-wise distributions. It models the probability of zero counts as a function of the mean expression of the gene and the library size (read depth) of the cell (both in log scale). Zero counts are then added to the simulated data such that the observed relationship (zero probability to mean expression and library size) is maintained. It simulates DE by separately estimating the distributions of the gene expression from the different populations (for example treatment groups) in the source data, and subsequently sampling a new dataset from each group. It is implemented using the *SPsimSeq* R package (version 2.0.0)[15].

### 2.6 Competitor tools for differential expression analysis

We compared the PIM method with six other tools for testing DE. We briefly discuss each method in the subsequent paragraphs.

**MAST** uses a hurdle model in which the zero counts are modelled by logistic regression, and the log TPM (or CPM) of the non-zero counts are modelled by normal linear regression [12]. Both models include the cellular detection rate (fraction of detected genes per cell) as a covariate. MAST is implemented using the *MAST* R biocondictor package (version 1.8.1)[12].

**edgeR** is a very common tool for testing DE in bulk RNA-seq data. It fits NB regression models with moderated estimation of the gene-specific over-dispersion parameters [30]. edgeR is implemented using the *edgeR* [20] R bioconductor package (version 3.22.5).

**DESeq2** relies on the NB distribution for testing DE in bulk RNA-seq data like edgeR. It uses an empirical Bayes moderation for estimating the over-dispersion parameters. It is implemented using the *DESeq2* R bioconductor package (version 1.20.0) [31].

**Zinger** calculates weights that can be used by edgeR and DESeq2 to account for the zero inflation in scRNA-seq data. In particular, Zinger fits zero-inflated negative binomial models for calculating weights that indicate the probability that the observed zero expression belongs to the NB component [10]. In this study, Zinger is used in conjunction with edgeR and DESeq2 (edgeR+Zinger and DESeq2+Zinger). It is implemented using the *Zinger* [10] R package (version 0.1.0).

**SAMSeq** is a non-parametric method which was originally developed for testing DE in bulk RNA-seq data. It uses the Wilcoxon statistic in conjunction with a permutation method to estimate the false discovery rate to infer DE [13]. It is implemented using the *samr* R software package (version 3.0) [32].

### 2.7 Real data analysis

We applied the seven DE tools for testing DE between nutlin-3 treated and control cells from two datasets. The datasets first went through pre-processing steps using the *scatter* R bioconductor package (version 1.6.3, [3]), which involves filtering cells with low quality metrics and genes with insufficient expression. The cell cycle stage is estimated using the *cyclone* function in the *scran* [33] R bioconductor package (version 1.10.1). In particular, we compared the number of detected DE genes at the 5% nominal FDR among the competitor tools. We also looked at the distribution of the log_2_(CPM+1) as well as the distribution of zero counts within the set of DE genes detected by each tool. We also explored the cross data agreement of all tools in detecting DE genes. In addition to DE testing, we performed gene set enrichment analysis (GSEA) using the *fgsea* R bioconductor package (version 1.8.0, [34]). In particular, we used a set of TP53 pathway genes obtained from [35]. Genes were ranked based on their test statistic as in [10].

### 2.8 Software implementation for PIM

An R software package for DE analysis using PIM is accessible from a GitHub repository with the name PIMseq (https://github.com/CenterForStatistics-UGent/PIMseq). Additionally, R codes used for all analyses reported in this manuscript are available in a GitHub repository (https://github.com/CenterForStatistics-UGent/PIMseq-paper).

## 3 RESULTS

### 3.1 Performance evaluation through simulation study

The ability to control the FDR without compromising the sensitivity to detect truly DE genes is an important requirement. This capability is evaluated in simulation studies in which gene expression data are generated with a built-in truth while retaining the characteristics of real data. Using three simulation procedures (Splat, SPsimSeq, and mock comparison), we evaluated the performance of PIM compared to edgeR, edgeR with Zinger, DESeq2, DESeq2 with Zinger, MAST and SAMSeq. The simulation methods vary with respect to the data generating model (see Methods section). In general, two sets of simulations were conducted, each starting from other source data (see Table 1). Each simulation set involves the three simulation methods. The performance of the DE methods is evaluated by calculating the actual FDR and true discovery rate (TPR) over the simulation runs.

We compared the simulated datasets with the real datasets with respect to various characteristics proposed by [9, 26], such as the distribution of mean and variance expression levels, mean-variance trend, distribution of zero fraction per gene, and variability among cells (Supplementary Figures S1 and S2). Although the Splat simulation is flexible in simulating various scenarios, the simulated data showed the least resemblance with the real datasets. The SPsimSeq procedure generated more realistic scRNA-seq data (Supplementary Figures S1 and S2).

Results from the simulation studies starting from source data A show that PIM succeeds in controlling the actual FDR below the nominal level (nominal FDR ranging from 0 to 0.3) for both Splat and SPsimSeq simulation procedures (Figure 2). This result is confirmed by simulations with different numbers of cells (Supplementary Figure S3) and with different magnitudes of the LFC of DE genes (Supplementary Figure S4). The actual TPR for PIM, SAMSeq and MAST are comparable and considerably higher (up to 20% increase) than the actual FDR for edgeR and DESeq2 (both with and without Zinger) for the SPsimSeq simulation when the LFC of DE genes is greater than 1 (Figure 2 and Supplementary Figure S3). For the SPsimSeq simulations, when the LFC of DE genes is low, PIM and SAMSeq outperform all the other tools with respect to TPR (Supplementary Figure S4). With the Splat simulation, edgeR and DESeq2 (both with and without Zinger) show better TPR (up to 10% increase) with the actual FDR below the nominal level. However, edgeR (with and without Zinger) loose control of the actual FDR for the SPsimSeq simulations, which do not make use of parametric distributional assumptions for the gene expression data (Figure 2 and Supplementary Figure S3). The good results for edgeR and DESeq2 are generally expected because the Splat method generates data from the negative binomial distribution, which is also the working model for both edgeR and DESeq2. However, the gap reduces to less than 5% when the number of cells per group is more than 100 (Supplementary Figure S3). In addition, for the Splat simulation, edgeR and DESeq2 show improved TPR when they are used with Zinger, especially when the LFC of DE genes is low (Supplementary Figure S4 and Figure 2).

**Figure 2:**
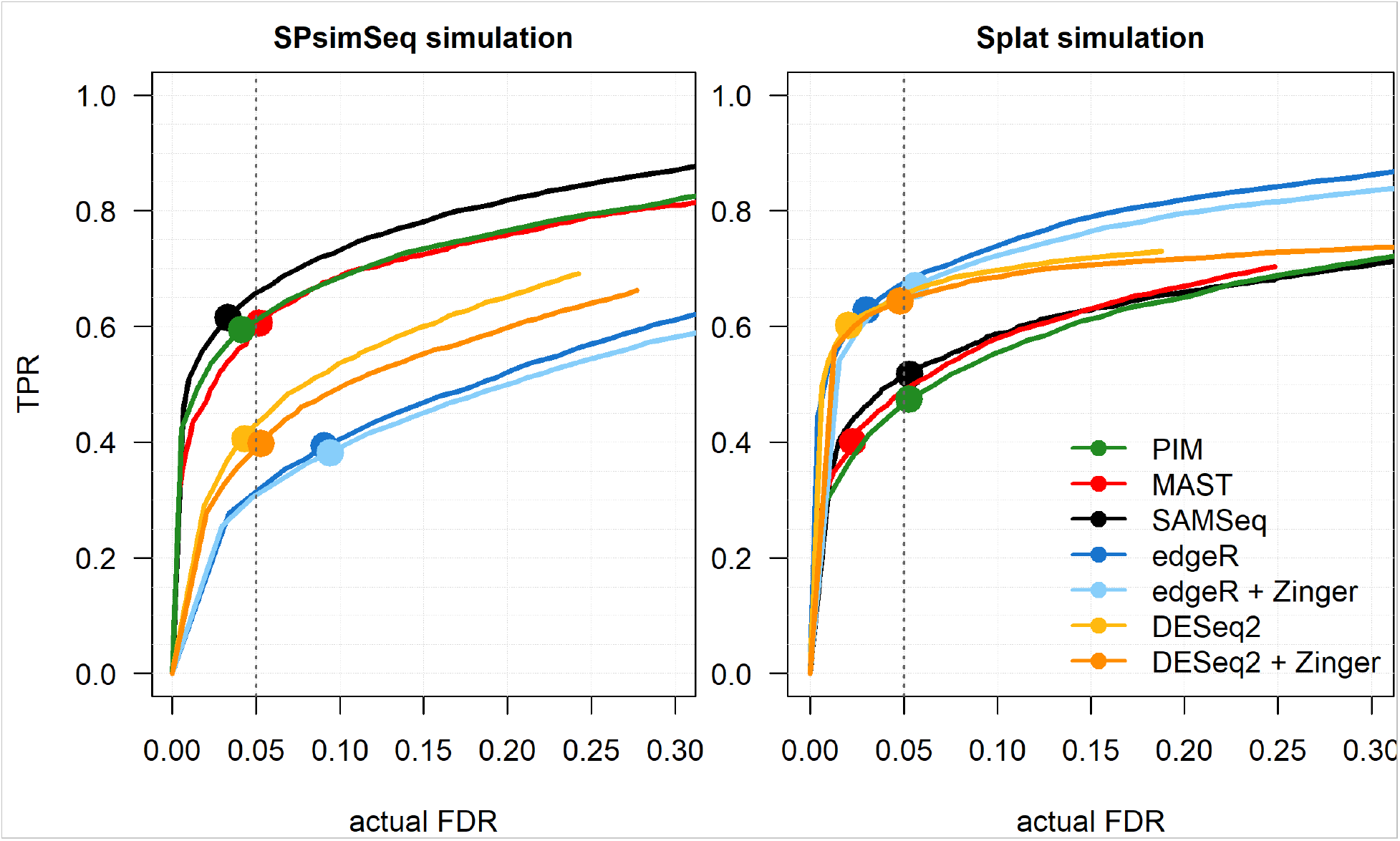
Results from the simulation study starting from source dataset A. Dataset A is neuroblastoma scRNA-seq data generated with the SMARTer/C1 protocol. Each simulated dataset includes 2500 genes among which 10% DE, and 2 experimental groups with each 50 cells. For the SPsimSeq simulation, the DE genes have LFC *≥* 1 in the source data, whereas for the Splat simulation the FC for DE genes is sampled from a log-Normal(location=1.5, scale=0.4), such that more than 97.5% of the DE genes have a LFC of at least 1. The performance measures (actual FDR and TPR) are averaged over a total of 50 independent simulation runs. The curves represent actual FDR and TPR evaluated at nominal FDR levels ranging from 0 to 0.3, and the solid dots show the performances at the 5% nominal FDR level.

The results starting from dataset B showed that all tools have rather small TPR, even though the number of cells is double of that of dataset A (Figure 3). This may be explained by the fact that scRNA-seq data with UMI-counts contain a large fraction of zero counts, low expression magnitude and large variability of the library sizes [11]. However, with LFC *≥* 1 for DE genes and 100 cells per group (Supplementary Figure S5) or with LFC *≥* 0.5 for DE genes and 200 cells per group (Supplementary Figure S6), all tools showed better performance than the performance in Figure 3 (for LFC *≥* 0.5 and 100 cells per group). With the SPsimSeq simulations, PIM shows better performance than all other tools for both simulation settings (Figure 3, Supplementary Figure S5 and S6). With the Splat simulations, PIM shows comparable performance to edgeR (with and without Zinger) and MAST when the LFC of DE genes is *≥* 0.5 (Figure 3 and Supplementary Figure S6). With the Splat simulations, when the LFC of DE genes is *≥* 1 PIM achieved the highest TPR while controlling the FDR slightly below the nominal level (Supplementary Figure S5). With the SPsimSeq simulations, SAMSeq and MAST and DESeq2 (with and without Zinger) generally show inferior performance, whereas with the Splat simulations, SAMSeq and DESeq2 (with and without Zinger) show relatively poor performance with respect to TPR (Figure 3, Supplementary Figure S5 and S6).

**Figure 3:**
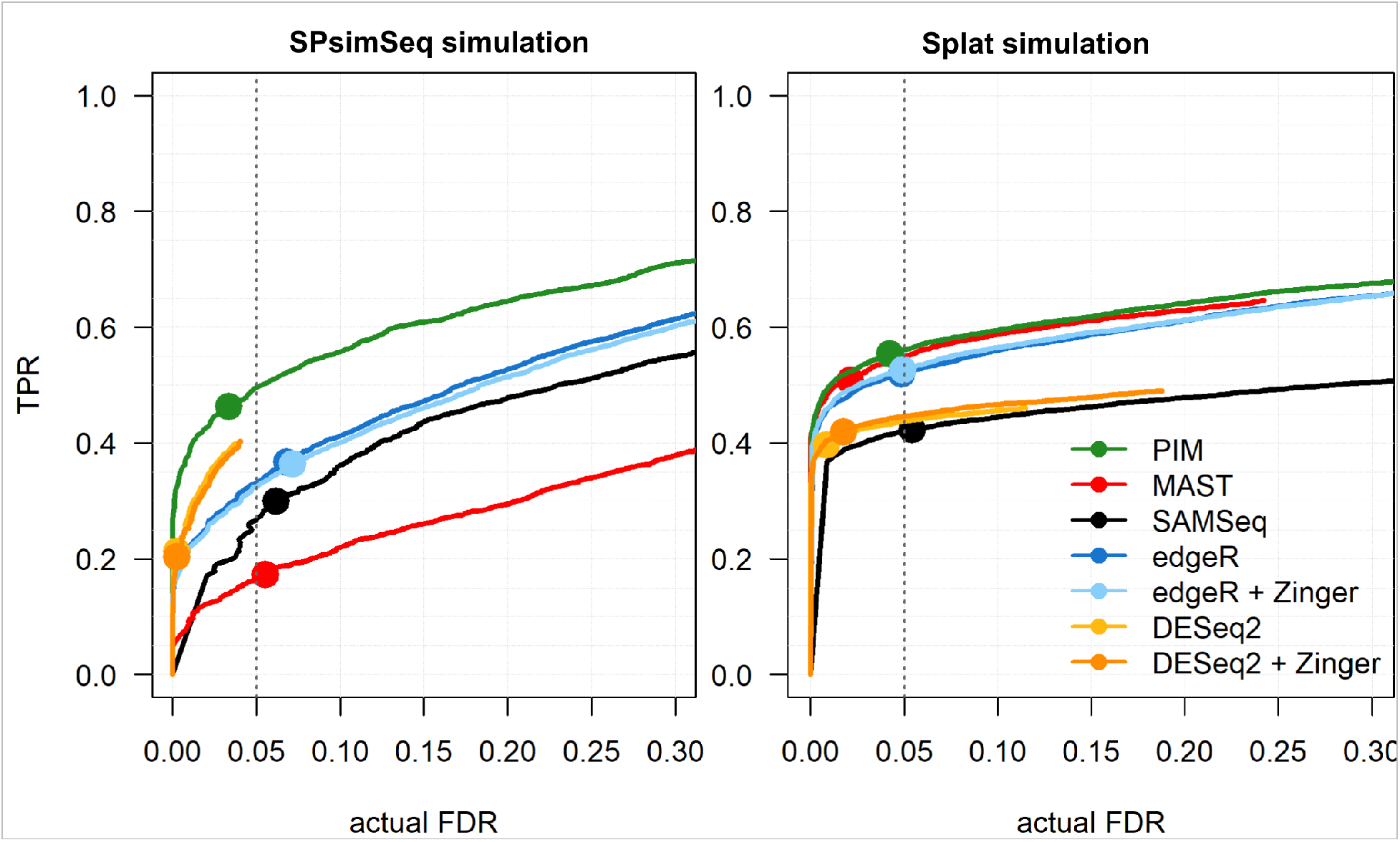
Results from the simulation study starting from source dataset B. Dataset B is neuroblastoma scRNA-seq data generated with the Chromium protocol (UMI counts). Each simulated dataset includes 2500 genes among which 10% DE, and 2 experimental groups with each 100 cells. For the SPsimSeq simulation, the DE genes have LFC *≥* 0.5 in the source data, whereas for the Splat simulation the FC for DE genes is sampled from a log-Normal(location=1.25, scale=0.4), such that more than 97.5% of the DE genes have a LFC of at least 0.5. The performance measures (actual FDR and TPR) are averaged over a total of 50 independent simulation runs. The curves represent actual FDR and TPR evaluated at nominal FDR levels ranging from 0 to 0.5, and the solid dots show the performances at the 5% nominal FDR level.

In order to demonstrate that PIMs can be applied to other settings than comparing gene expression between two groups, we simulated a single cell RNA-seq data with three biological groups. In this setting, PIM remains applicable along with edgeR, DESeq2 and MAST, with comparable performance (Supplementary Figure S7).

We also evaluated the capability of PIM in controlling the type-I error rate using mock comparisons. Since SAMSeq does not return un-adjusted p-values, it is excluded from this comparison. Results from the mock comparison are presented in two ways. First, the observed average false positive proportions (FPP) is nearly equal to the nominal 5% per-comparison error rate (PCER) for all tools (Supplementary Figure S8). As the subsequent results show, the parametric tools are relatively conservative, leading to smaller numbers of false positives. Second, since mock comparison simulates gene expression data under the null hypothesis (no DE), the distribution of raw p-values is expected to be uniform between 0 and 1. The distribution of raw p-values for PIM and all other considered DE tools are uniform except for DESeq2 (and to same degree also not for MAST), which shows a substantial deviation with higher probability mass near p-value=1 (also known as a conservative distribution; Figure S8). This result is in line with the smallest observed average FPP for DESeq2 compared to all other tools (Supplementary Figure S9). For uniform p-value distributions, as for PIM, it is anticipated that FDR control procedures, such as Benjamini–Hochberg (BH) [19], can be used for FDR control.

### 3.2 DE analysis of nutlin-3 treatment of neuroblastoma cells

We applied PIM and the competitor tools for testing DE in two nutlin-3 treated neuroblastoma scRNA-seq datasets. Nutlin-3 is a selective MDM2 antagonist, releasing TP53 from its targeted degradation, resulting in cell cycle arrest, followed by apoptotic cell death. The objective is to identify genes with DE between nutlin-3 treated and vehicle (control) cells. We focus on (i) comparing the number of genes detected as DE (at 5% nominal FDR) between PIM and other tools, (ii) exploring the set of DE genes in terms of the distribution of gene expression and fold change estimates (PI for PIM), (iii) TP53 gene set enrichment score, and (iv) cross-data agreement.

For dataset A, we fitted the PIM with the treatment group and the log library size in the model. In this way, we test for DE by accounting for the library size differences. For dataset B, we extended the PIM model by adding the cell cycle stage (a discrete factor with three levels for the three cell cycle stages: G1, G2/M, and S) in the model. From dataset A, PIM identified a set of 355 genes, 225 down-regulated and 130 up-regulated genes in the nutlin-3 treated group (Figure 4–5). Compared to the other tools, PIM ranks second following SAMSeq, which detected over 430 DE genes. The parametric tools MAST, edgeR and DESeq2 identified relatively small numbers of DE genes (Figure 4). PIM and SAMSeq identified 96 DE genes in common (Figure 4a).

**Figure 4:**
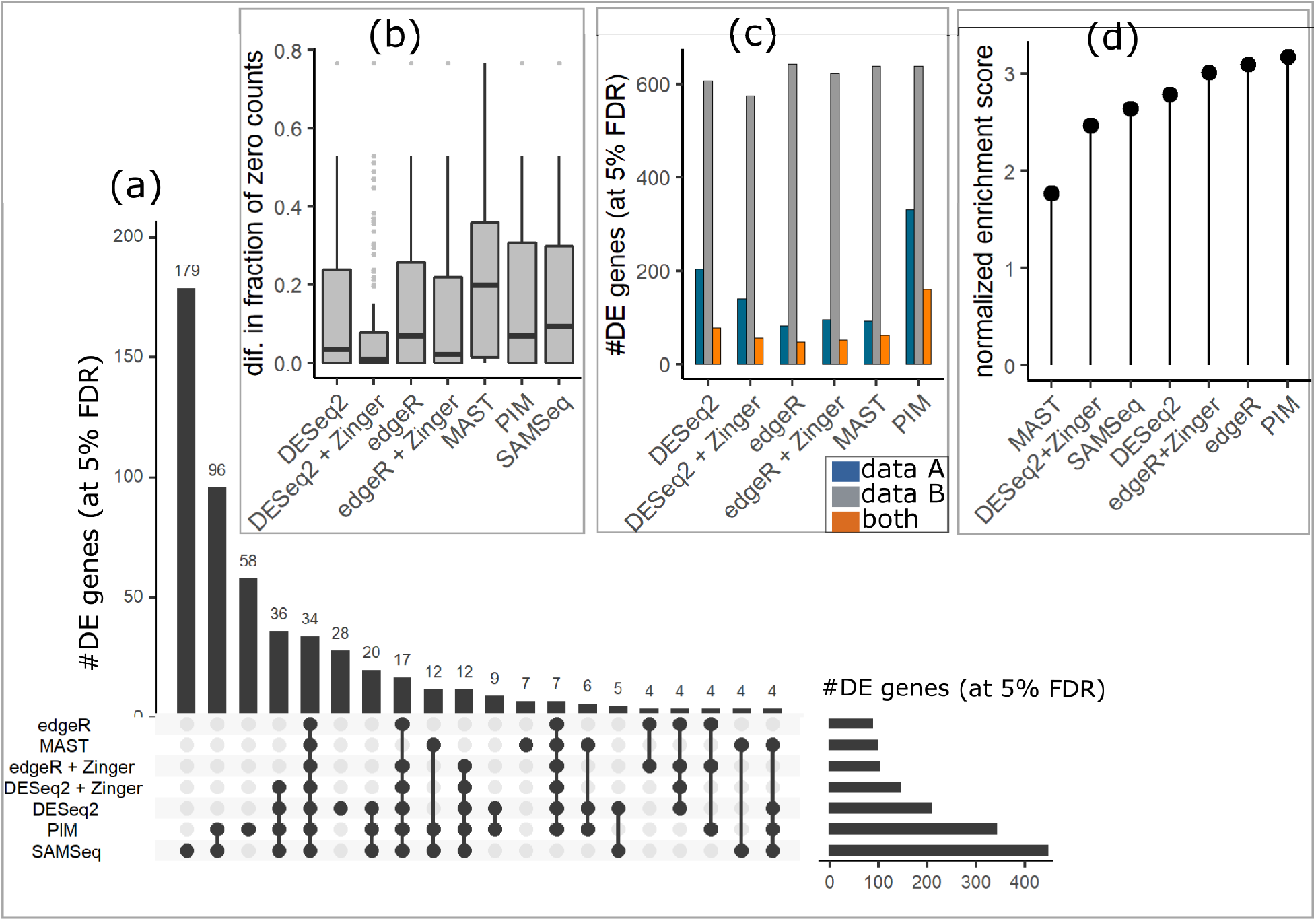
Results from DE analysis of dataset A and B. (a) UpSet plot to compare the set of DE genes detected by the 7 tools at the 5% FDR level. The vertical bars show the number of genes called DE by all tools indicated by the connected black dots (intersection), whereas the row bars indicate the number of genes called DE by each tool at the 5% FDR level, (b) the distribution of the difference (absolute) in the fraction of zero counts between nutlin-3 treated and control groups, for the set of genes called DE by each tool, (c) the number of genes called DE by each tool in the two data sets (datasets A and B) and the number of DE genes in both datasets by each tool (cross data agreement), and (d) the normalized enrichment score for TP53 gene set.

SAMSeq and PIM called the highest number of DE genes from dataset A at the 5% FDR level (Figure 4a). In particular, these tools identified almost all the DE genes identified by all the parametric tools. From exploring the distribution of the gene expressions (log-CPM), specifically for the set of DE genes, we learn that SAMSeq and MAST tend to pick genes with a large difference in the fraction of zero counts between the two treatment groups (Figure 4b, Supplementary Figure S10). In contrast, the set of DE genes identified by edgeR+Zinger and DESeq2+Zinger have the smallest difference in the distribution of zero counts (also compared to edgeR and DESeq2 without Zinger), presumably because Zinger down-weights the contribution of zero counts (Figure 4b). Consequently, edgeR+Zinger and DESeq2+Zinger mostly identify genes with difference only in the distribution of the non-zero expressions (Supplementary Figure S10). PIM detected DE genes with small to moderate differences in zero count distributions across the groups (Figure 4b, Supplementary Figure S11). SAMSeq uniquely detected almost 180 DE genes with the majority of them having a high level of variability (highly expressed genes) and small LFC estimate (*<* 0.5) (Supplementary Figure S11). Soneson and Robinson [9] have also noticed that SAMSeq is biased towards highly expressed genes and genes with a small number of zero counts. A similar set of DE genes are commonly detected by both PIM and SAMSeq (Supplementary Figure S11). However, among the DE genes uniquely identified by PIM, the number of highly variable genes and genes with *<* 0.5 LFC is relatively small (Supplementary Figure S11). In the simulation study starting from dataset A, PIM, SAMSeq, and MAST showed similar performance when the LFC of DE genes is at least 1 (Figure 2). However, when the LFC of DE genes is at least 0.5, PIM and SAMSeq perform similarly and better than all other tools (Supplementary Figure S4). This result is in line with the empirical result that PIM and SAMSeq have the capability of detecting DE genes with small to high LFCs.

For dataset B, all tools but SAMSeq called many genes DE at the 5% FDR level because of the large number of cells in dataset B (Figure 4c). The two datasets are similar in terms of experimental design, but they resulted from different protocols (SMARTer/C1 and Chromium) and contain different gene expression units (read counts and UMI counts). In both datasets, we test for DE between the nutlin-3 and control group. This further allows us to evaluate the cross-data agreement of the tools. SAMSeq failed for dataset B and was excluded from this assessment. PIM identified 152 common DE genes in the two datasets, whereas MAST, edgeR, edgeR+Zinger, DESeq2 and DESeq2+Zinger identified 62, 48, 51, 78, and 52 genes, respectively, in both datasets (Figure 4c). This result suggests that PIM has better consistency across protocols and gene expression units. SAMSeq fails with no particular error message during the re-sampling step. This issue has also occurred in the simulation studies starting from dataset B (Supplementary Figure S6).

The model parameters of PIM have an informative interpretation as an effect size in terms of the PI. We now demonstrate that the estimated PI is a useful metric in addition to the fold-change used in parametric DE analysis. Fold changes express the biological signal in terms of the difference in the mean expression between conditions, with unbounded scale. Similarly, the PI quantifies the degree of difference in the distribution of expression at a scale ranging from 0 to 1 (see Figure 1). In Figure 5a, we ranked genes according to their estimated PI in such a way that genes with strong evidence of DE appear near the left edge (if down-regulated in treated group) or near the right edge (if up-regulated in treated group). The estimated PIs are linearly correlated with the LFC (Figure 5b). Note that we do not suggest that the LFC should not be used. With parametric methods, the fold-change is still the effect size that correspond to the p-value. In a typical gene expression study, the majority of the genes are not DE, with estimated LFC close to 0. This also holds for PIM, i.e. the estimated PI for the majority of the genes is around 0.5 (Figure 5a and reffig:Fig5c).

**Figure 5:**
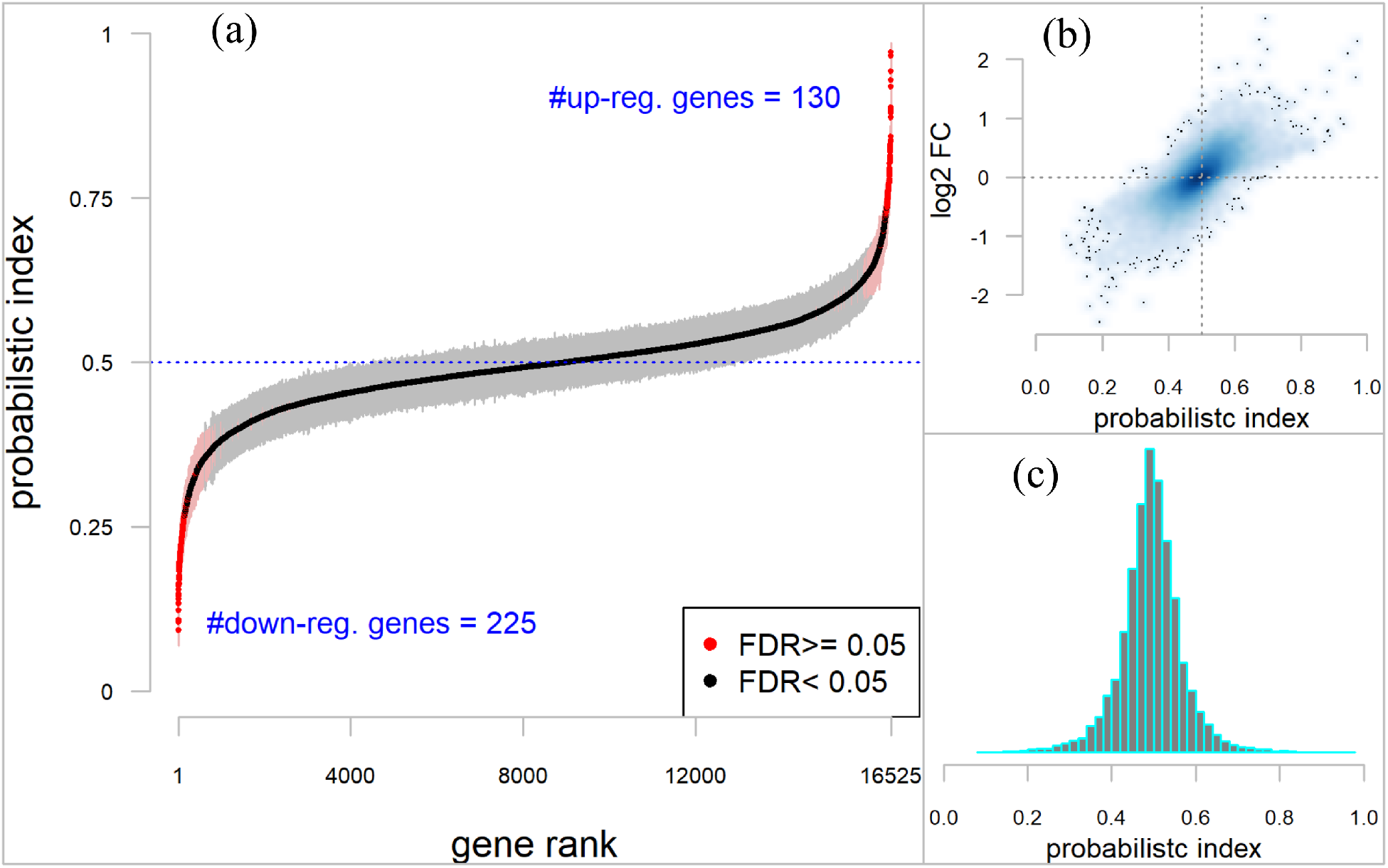
PIM results for the DE analysis of dataset A. (a) Gene ranking based on their estimated PI. The gray region indicates the 95% confidence interval estimate of the PI for each gene. (b) The relationship between estimated PI and a log_2_-fold-change (log_2_FC) estimates. The log_2_FC were obtained using DESeq2. (c) The distribution of PI across genes.

We also ran GSEA to find out if the DE genes detected by PIM may have a biological meaning. Nutlin-3 liberates TP53 from MDM2-mediated inhibition and hence activates the TP53 transcriptional response pathway [35]. Therefore, we used a set of 116 TP53 pathway consensus genes [35]. We compared the normalized enrichment score (NES) among the 7 DE tools including PIM (Figure 4c and Supplementary Figure S12). All the tools scored above 1.7 NES with p-value *<* 0.001, and PIM attained the highest NES (NES=3.16, p-value 0.00044) followed by edgeR (NES=3.11, p-value 0.00035). Generally, these results indicate that PIM showed significant enrichment of the TP53 pathway in neuroblastoma cells treated with nutlin-3.

### 3.3 Computation time

The computation time of PIM in comparison to the other tools is evaluated using five simulated scRNA-seq datasets each with a different number of cells, ranging from 100 to 10,000 (Table 2). All the datasets contain 10,000 genes. The computation time for PIM increases quadratically with the number of cells. For example, for 200 cells, PIM requires less than 2 minutes, whereas for 1000 cells, it requires up to 1 hour (Table 2). However, with the Cox-PH model translation of the PIM, it requires less than 3 minutes for 10,000 cells. MAST and edgeR are among the fastest methods, and DESeq requires more than 5 hours for 2000 cells. Zinger is generally computationally intensive, especially when coupled with DESeq2.

**Table 2:** Computation time in minutes. All tools were implemented on a single core CPU (Core i7-6820HQ and 16.0 GB RAM) with Windows 10 operating system.

| tool | number of cells |  |  |  |  |
| --- | --- | --- | --- | --- | --- |
|  | 100 | 200 | 1000 | 2000 | 10,000 |
| PIM | 0.84 | 1.63 | 62.53 | > 5h | > 5h |
| PIM (Cox-PH) | 0.37 | 0.38 | 0.55 | 0.76 | 2.57 |
| SAMSeq | 0.19 | 0.34 | – | – | – |
| MAST | 0.92 | 0.89 | 1.24 | 1.53 | 4.03 |
| edgeR | 0.14 | 0.26 | 1.18 | 2.37 | 12.67 |
| edgeR+Zinger | 3.63 | 4.44 | 21.50 | 42.45 | > 5h |
| DESeq2 | 0.28 | 0.69 | 30.52 | > 5h | > 5h |
| DESeq2+Zinger | 3.06 | 4.33 | 71.83 | > 5h | > 5h |

> 5h= more than 5 hours
– SAMSeq fails for datasets with > 200 cells

## 4 DISCUSSION

Advances in sequencing technology enable profiling of gene expression levels at individual cell level. ScRNA-seq experiments are used for various applications in the field of biology and medicine, such as identifying genes whose expression differ under different conditions (also known as DE). Importantly, scRNA-seq data are characterized by high levels of noise resulting from various technical sources, such as limited RNA capture efficiency, amplification bias and differences in RNA content [1, 5]. Furthermore, the diverse biological characteristics of individual cells and the stochastic nature of gene expression challenge computational tools for testing DE.

Two broad classes of methods exist for testing DE, differing in terms of their underlying assumption for the gene expression data, i.e. parametric and non-parametric. Parametric tools such as edgeR and DESeq2 have been widely used for bulk and single cell RNA-seq studies. When used in conjunction with Zinger [10], they account for both the over-dispersion and zero-inflation issues in scRNA-seq data. MAST fits hurdle-Gaussian models to deal with zero inflation assuming that all zeroes result from technical factors. The performance of these tools seemed adequate using simulation studies that make use of their underlying distribution [9]. In contrast, non-parametric methods require minimal distributional assumptions and are considered useful when the underlying data generating distribution is unknown, as is the case for scRNA-seq data. Simple non-parametric methods like the Wilcoxon rank-sum test shows competitive or better performance relative to many of the parametric tools specifically developed for scRNA-seq data [9, 14]. Unfortunately, the usual non-parametric tools have a limited utility for scRNA-seq studies with complex experimental design. Moreover, they do not provide an informative estimate of the effect size, which is a vital component of DE analysis. In this paper we have introduced a semi-parametric method for testing DE in scRNA-seq data based on PIMs, which generalize many of the non-parametric methods and provides an informative effect size measure.

Using three simulation studies, we have demonstrated that PIMs succeed in controlling the FDR and that it has reasonable sensitivity to detect true DE genes. The three simulation studies differ with respect to the underlying data generating mechanism: fully parametric, semi-parametric, and non-parametric (mock comparison). Unlike the parametric tools, PIMs performs well under both simulation mechanisms for various magnitudes of the biological effect (LFC) and small to large numbers of cells. In contrast, edgeR and DESeq2 showed slightly better performance when the data were generated from negative binomial distributions. MAST and SAMSeq are greatly affected by the high fraction of zeroes in simulation studies involving UMI counts. We also applied PIMs and the other tools to two real and novel scRNA-seq datasets. These datasets represent two different types of experiments with different platforms, numbers of cells and ‘units’ of gene expression, i.e. raw read counts versus UMI deduplicated reads. The results demonstrate that PIMs detect large numbers of DE genes with better cross-data agreement. Further, the PI ranked gene list is significantly positively enriched with genes from a pathway known to be activated in the cellular model system, demonstrating that PIM allows to attain biological insights from scRNA-seq experiments.

We have also demonstrated that PIMs can be scalable and computational efficient for experiments with large number of cells with the Cox-proportional hazard model translation of PIMs. Finally, while we demonstrated the utility of PIMs for testing DE in scRNA-seq data, PIMs could also be used for bulk RNA-seq and microarray datasets with a sufficient number of replicates. In principle, PIMs can also be integrated with other data pre-processing pipelines such as normalization, batch correction, and imputation, making it a useful and general tool for single cell RNA seq differential gene expression analysis.

## SUPPLEMENTARY DATA

Additional supporting results and discussion are available in *Supplementary File*.

## Author’s contributions

ATA: Conceptualization, Data curation, Formal analysis, Methodology, Software, Visualization, Writing – original draft, Writing – review & editing. JV: Conceptualization, Funding acquisition, Methodology, Supervision, Writing – review & editing. OT: Conceptualization, Funding acquisition, Methodology, Supervision, Software, Writing – review & editing.

