## supplementary file for "Probabilistic index models for testing differential expression in single cell RNA sequencing data"

*February 20, 2020*

**3 Additional results from analysis of real scRNA-seq datasets** **14**

**References** **16**

#### 1 PIM parameter estimation and further properties

Let $Y_{gi}$ be the gene expression of gene $g = 1, 2, \ldots G$ in cell $i = 1, 2, \ldots, n$ and let $\mathbf{X}_i$ is a $p \geq 1$ dimensional vector with cell level covariates. For $n$ i.i.d. sample observation $(Y_{gi}, \mathbf{X}_i)$, $i = 1, \ldots, n$, a PIM models the conditional probability

$$\mathrm{P}(Y_{gi} \preccurlyeq Y_{gj}|\mathbf{X}_i, \mathbf{X}_j) = \mathrm{P}(Y_{gi} < Y_{gj}|\mathbf{X}_i, \mathbf{X}_j) + \frac{1}{2}\mathrm{P}(Y_{gi} = Y_{gj}|\mathbf{X}_i, \mathbf{X}_j), \tag{1}$$

[1] defined the PIM as

$$\mathrm{P}(Y_{gi} \preccurlyeq Y_{gj}|\mathbf{X}_i, \mathbf{X}_j) = m(\mathbf{X}_i, \mathbf{X}_j; \boldsymbol{\beta}_g) = g^{-1}(\mathbf{Z}_{ij}^T \boldsymbol{\beta}_g), \tag{2}$$

where $\mathbf{Z}_{ij} = \mathbf{X}_j - \mathbf{X}_i$, $g(.)$ is a link function, $m(.)$ is a function that has a range [0, 1], and $\boldsymbol{\beta}_g \in \mathbb{R}^p$ is a $p$ dimensional vector of parameters. This model is defined for a set of pairs of regressors $(\mathbf{X}_i, \mathbf{X}_j)$, which is denoted as $\chi_n$. Similar to a GLM, the systematic component $m(\mathbf{X}_i, \mathbf{X}_j; \boldsymbol{\beta}_g)$ is restricted to $g^{-1}(\mathbf{Z}_{ij}^T \boldsymbol{\beta}_g)$ with an appropriate link function $g(.)$, such that

- $m(., .; \boldsymbol{\beta}_g)$ has a range [0, 1]
- if $(\mathbf{X}_i, \mathbf{X}_i)$, $(\mathbf{X}_i, \mathbf{X}_j)$, and $(\mathbf{X}_j, \mathbf{X}_j)$ are elements of $\chi_n$, then $m(., .; \boldsymbol{\beta}_g)$ must satisfy
  - $m(\mathbf{X}_i, \mathbf{X}_i; \boldsymbol{\beta}_g) = 0.5$ and
  - $m(\mathbf{X}_i, \mathbf{X}_j; \boldsymbol{\beta}_g) + m(\mathbf{X}_j, \mathbf{X}_i; \boldsymbol{\beta}_g) = 1$.

The choice $\mathbf{Z}_{ij} = \mathbf{X}_j - \mathbf{X}_i$ and the logit link function $(g(x) = x/(1 - x))$ provide an simple interpretation of the PIM.

For the above model definition with few more assumptions, [1] proposed a semi-parametric consistent and asymptotically normally distributed estimator of $\boldsymbol{\beta_g}$. In particular, the estimation functions use a set of pseudo-observations defined as

$$I_{ij(g)} = \mathrm{I}(Y_{gi} \preccurlyeq Y_{gj}) := \mathrm{I}(Y_{gi} < Y_{gj}) + 0.5\mathrm{I}(Y_{gi} = Y_{gj}) \quad \text{for all } i,\ j = 1, 2, \ldots, n \text{ for which } \boldsymbol{X_i}, \boldsymbol{X_i} \in \chi_n$$

where I(.) is the usual 0/1 indicator function. The motivation behind the pseudo-observations $I_{ij(g)}$ is that the probabilistic index is the expectation of $I_{ij(g)}$, i.e.

$$P(Y_{gi} \preccurlyeq Y_{gj}|\mathbf{X}_i, \mathbf{X}_j) = \mathrm{E}\left\{\mathrm{I}(Y_{gi} \preccurlyeq Y_{gj}|\mathbf{X}_i, \mathbf{X}_j)\right\}$$

.

Thus $\boldsymbol{\beta}_g$ can be estimated by solving the estimating equations

$$\mathbf{U}_{n(g)}(\boldsymbol{\beta}_g) = \sum_{i,j \in \daleth_n} \mathbf{U}_{ij(g)}(\boldsymbol{\beta}_g) = \mathbf{0} \tag{3}$$

with

$$\mathbf{U}_{ij(g)}(\boldsymbol{\beta}_g) = \mathbf{A}(\mathbf{Z}_{ij}, \boldsymbol{\beta}_g)\left\{I_{ij(g)} - g^{-1}(\mathbf{Z}_{ij}^T\boldsymbol{\beta}_g)\right\},$$

and $\daleth_n$ is the set of indices $I_{ij(g)}$ for which $\mathbf{X}_i,\ \mathbf{X}_j \in \chi_n$, and $\mathbf{A}(\mathbf{Z}_{ij}, \boldsymbol{\beta}_g)$ is a $p$-dimensional vector-valued function of $\mathbf{Z}_{ij}$ given by

$$\mathbf{A}(\mathbf{Z}_{ij}, \boldsymbol{\beta}_g) = \frac{\partial g^{-1}(\mathbf{Z}_{ij}^T\boldsymbol{\beta}_g)}{\partial \boldsymbol{\beta}_g}\mathbf{V}^{-1}\left\{g^{-1}(\mathbf{Z}_{ij}^T\boldsymbol{\beta}_g)\right\}$$

.

where $\mathbf{V}^{-1}\left\{g^{-1}(\mathbf{Z}_{ij(g)}^T\boldsymbol{\beta}_g)\right\} = (1/v)\mathrm{Var}(I_{ij(g)}|\mathbf{Z}_{ij})$, with $v$ a scale parameter.

[1] also demonstrated that $\sqrt{n}(\hat{\boldsymbol{\beta}}_g - \boldsymbol{\beta}_g)$ is asymptotically normally distributed with mean $\mathbf{0}$ and covariance matrix $\boldsymbol{\Sigma}_g$. Since the pseudo-observations $I_{ij(g)}$ are functions of pairs of the outcome variables $Y_{gi}$ and $Y_{gj}$, the $I_{ij(g)}$ are not longer mutually independent. Consequently, the usual covariance estimation methods that make use of the i.i.d. assumption are not appropriate in this scenario. As a result, [1] suggested a consistent sandwich estimator of $\boldsymbol{\Sigma}_g$, which is given by

$$\hat{\boldsymbol{\Sigma}}_g = \left(\sum_{i,j \in \daleth_n} \frac{\partial \mathbf{U}_{ij(g)}(\hat{\boldsymbol{\beta}}_g)}{\partial \boldsymbol{\beta}_g^T}\right)^{-1} \times \left(\sum_{i,j \in \daleth_n}\sum_{k,l \in \daleth_n} \phi_{ijkl(g)}\mathbf{U}_{ij(g)}(\hat{\boldsymbol{\beta}}_g)\mathbf{U}_{kl(g)}^T(\hat{\boldsymbol{\beta}}_g)\right) \times \left(\sum_{i,j \in \daleth_n} \frac{\partial \mathbf{U}_{ij(g)}(\hat{\boldsymbol{\beta}}_g)}{\partial \boldsymbol{\beta}_g^T}\right)^{-T} \tag{4}$$

where $\phi_{ijkl(g)} = 1$ if $\mathrm{I}(Y_{gi} \preccurlyeq Y_{gj})$ and $\mathrm{I}(Y_{gk} \preccurlyeq Y_{gl})$ are correlated and $\phi_{ijkl(g)} = 0$ otherwise.

The method is implemented in the R software package *pim* [2]. We use this package to further build an environment in which PIM can be used for testing DGE in single cell RNA-seq data. This is available as R software package named *PIMseq*, and it can be accessed from https://github.com/CenterForStatistics-UGent/PIMseq.

#### 1.1 Cox-Proportional hazard (Cox-PH) approximation of PIM

The parameter estimation methods for PIM involve comparison of outcome pairs, and thus the computation time increases quadratically with the sample size. As a result, for sequencing experiments with a large number of cells (such as > 500), we propose the Cox-proportional hazard [3] approximation of PIMs for computational efficiency.

[1], [4], and [5] have demonstrated that the Cox-PH model implies a PIM with logit link function and $\mathbf{Z}_{ij} = \mathbf{X}_j - \mathbf{X}_i$. In particular, we treat $Y_{gi}$ as the time to event for the $n$ independent cells and with

no censoring. Briefly, under the Cox-PH framework, for gene $g$ we model the hazard rate as $\lambda(y_g|\mathbf{X}) = \lambda_0(y)\exp(\mathbf{X}^T\boldsymbol{\beta}_g)$, where $\lambda_0(y_g) = \lambda(y_g|\mathbf{X} = 0)$ is the baseline hazard rate. From this model we can see that $\lambda(y_g|\mathbf{X})/\lambda_0(y_g) = \exp(\mathbf{X}^T\boldsymbol{\beta}_g)$, which is independent of $y_g$. This is also called the proportional hazard assumption.

The hazard rate $\lambda(y_g|\mathbf{X})$ is defined as $-\frac{\partial \log S(y_g)}{\partial y_g} = \frac{f(y_g)}{S(y_g)}$, where $f(y_g)$ is the density function of $y_g$ and $S(y_g)$ the corresponding survival function. The conditional survival function $S(y|\mathbf{X}) \propto S_0(y)^{\exp(\mathbf{X}^T\boldsymbol{\beta})}$, where $S_0(y_g) = S(y_g|\mathbf{X} = \mathbf{0})$ is the baseline survival function. From this result, it follows that

$$\mathrm{P}(Y_{gi} \preccurlyeq Y_{gj}|\mathbf{X}_i, \mathbf{X}_j) = -\int_S (1 - S(y_{gi}|\mathbf{X}_i))dS(y_{gj}|\mathbf{X}_j) = 1 - \exp\{\boldsymbol{\beta}_g(\mathbf{X}_j - \mathbf{X}_i)\}\mathrm{P}(Y_{gi} \preccurlyeq Y_{gj}|\mathbf{X}_i, \mathbf{X}_j),$$

which leads to the standard PIM defined above.

Parameter estimation of the Cox-PH model is based on maximizing the partial-log-likelihood function [6]. This estimation method is based on a restrictive assumption that survival time is continuous. However, scRNA-seq data consists of discrete observations, and hence it is likely that ties occur. Several alternative parameter estimation methods have been proposed to tackle this issue, and among these, researches showed that Efron's method[7] can be a better choice[8]. For a given gene $g$, we used Efron's approach that maximizes the following partial log-likelihood

$$\ell_g(\boldsymbol{\beta}_g) = \sum_j \left( \sum_{i \in H_{gj}} X_i \beta_g - \sum_{l=0}^{m_g - 1} \log \left( \sum_{i: Y_i \geq t_{gj}} \theta_{ig} - \frac{l}{m_g} \sum_{i \in H_{gj}} \theta_{ig} \right) \right),$$

where $\theta_{ig} = e^{X_j \beta_g}$, $t_{gj}$ is the unique times points (read counts), $H_{gj}$ is the set of indices $i$ such that $Y_i = t_j$ and $m_g = |H_{gj}|$. This is implemented by using the *coxph()* function of the *survival* R CRAN package [9]. Because this algorithm is computationally more efficient than that of the standard PIM estimation method discussed above, we have used it to estimate the PIM parameters. In particular, we use this estimation method for single cell data with large numbers of cells, especially for droplet-based protocols.

Although CoxPH implies PIM, the reverse is not necessarily true, because the proportional hazard assumption is not implied by the PIM. Therefore, we empirically examined the PH assumption using real single cell RNA-seq data. In particular, we used the two neuroblastoma cell line scRNA-seq datasets (datasets A and B; see Table 1 in main text), and we applied the Cox-PH model independently for each gene, with the treatment and the library size (in log-scale) as covariates. Afterwards, using the Schoenfeld residuals we assessed the PH assumption for both covariates and at a global level. This results in a vector of p-values for the three tests (for 2 covariates and global). Upon using the *fdrtool* R CRAN package [10] we estimated the frequency that the PH assumption was true. The results for the two datasets are summarized in Table S1. The results indicate that the PH assumption holds for at least 91% of the genes in scRNA-seq data with UMI-counts, and for 82.5% of genes in the data with read-counts. In general, the empirical results suggest that the PH assumption holds for majority of the genes and we therefore use the Cox-PH approximation of PIM. Note that PIM is less restrictive than the Cox-PH model, and results from PIM are valid when the PH assumption does not hold [5].

Table S1: Estimated fractions of genes for which the null hypotheses of the proportional hazard assumption holds true. The estimated fractions (standard error) are obtained with the *fdrtool* R CRAN package.

| | UMI-counts | read-counts |
|---|---|---|
| treatment | 0.990 (0.0022) | 0.991 (0.0028) |
| library size (in log) | 0.919 (0.0094) | 0.817 (0.0087) |
| GLOBAL* | 0.915 (0.0095) | 0.825 (0.0102) |

* a global test for all the factors together

### 2 Additional simulation results

#### 2.1 Simulations

In this section we show results of an empirical comparison between the real data and simulated data (Splat [11] and SPsimSeq [12] methods). In particular, we look at the distribution of the fraction of zero counts per gene, cell to cell similarity, the relationship between log mean expression and fraction of zero counts, and the relationship between log mean expression and coefficients of variation.

In this supplementary file, we show only a subset of the results. More results are available in [12]. On the following pages, we show the comparison for the two simulations: scRNA-seq data with read-counts (source data from SMARTer/C1 protocol = dataset A), and UMI-counts (generated using Chromium protocol = dataset B).

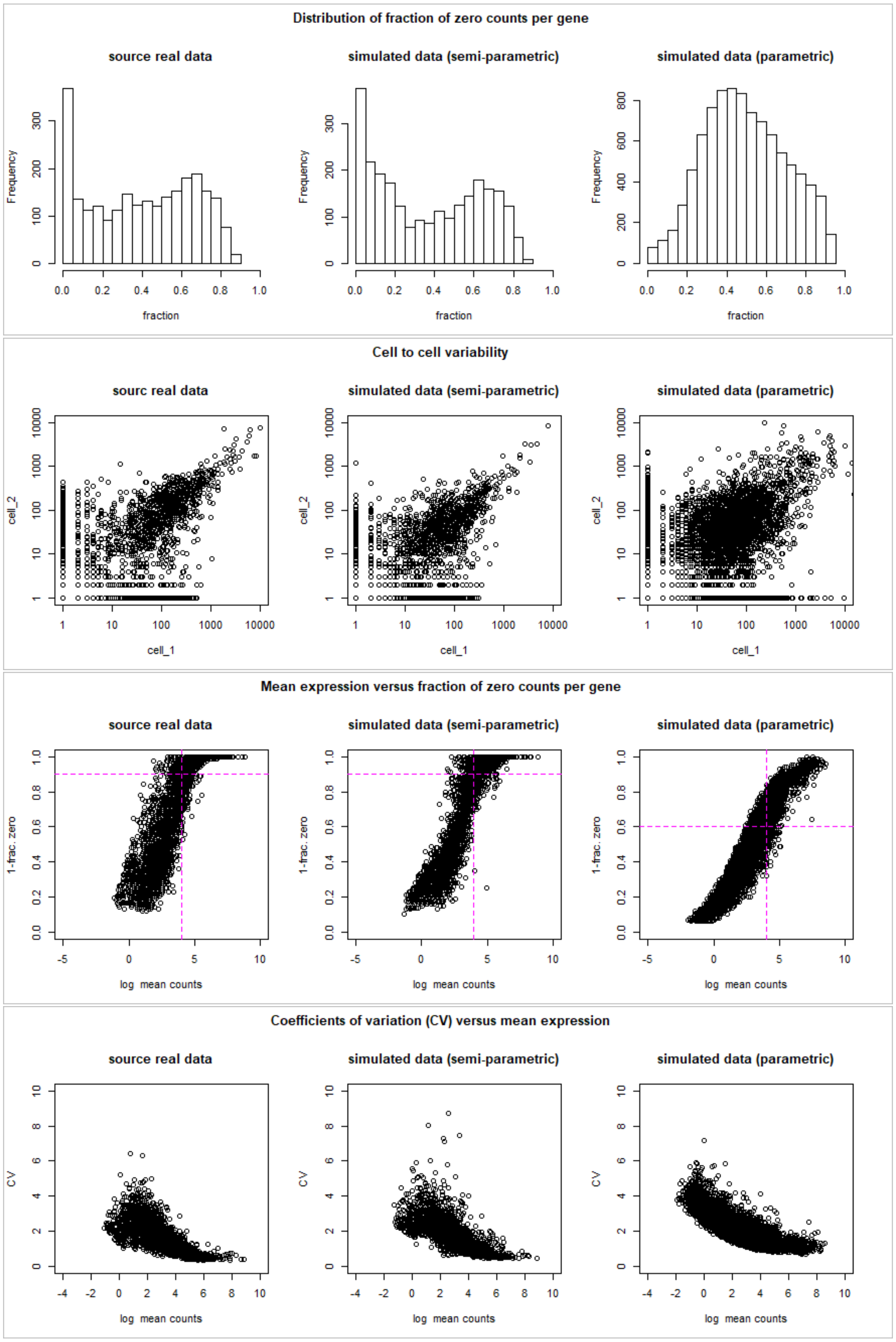

Figure S1: Comparison of the real data (source dataset A) and simulated data starting from this source data. The semi-parametric simulation implemented using SPsimSeq R software package and the parametric simulation is using the splatter R bioconductor package.

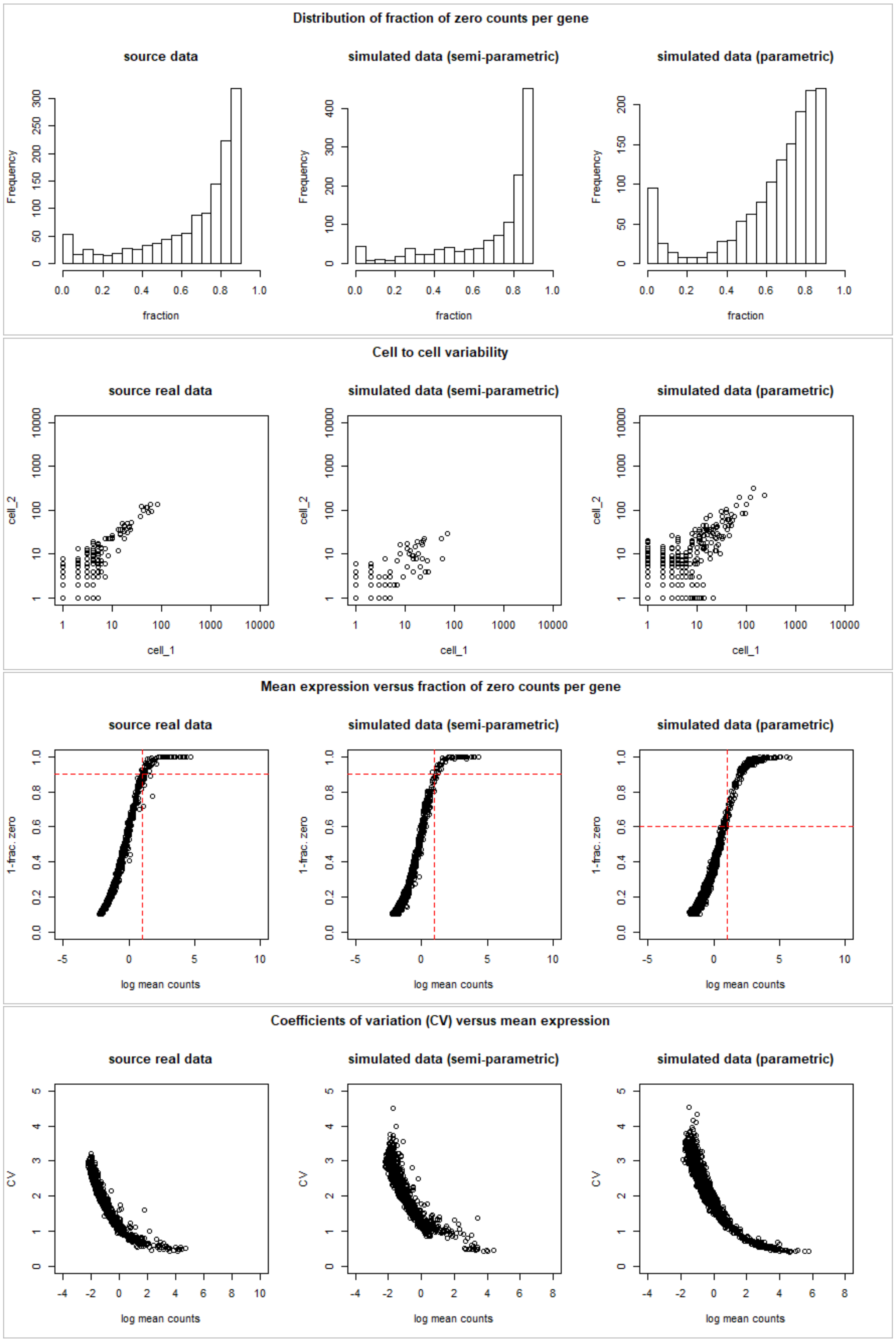

Figure S2: Comparison of the real data (source dataset B) and simulated data starting from this source data. The semi-parametric simulation implemented using SPsimSeq R software package and the parametric simulation is using the splatter R bioconductor package.

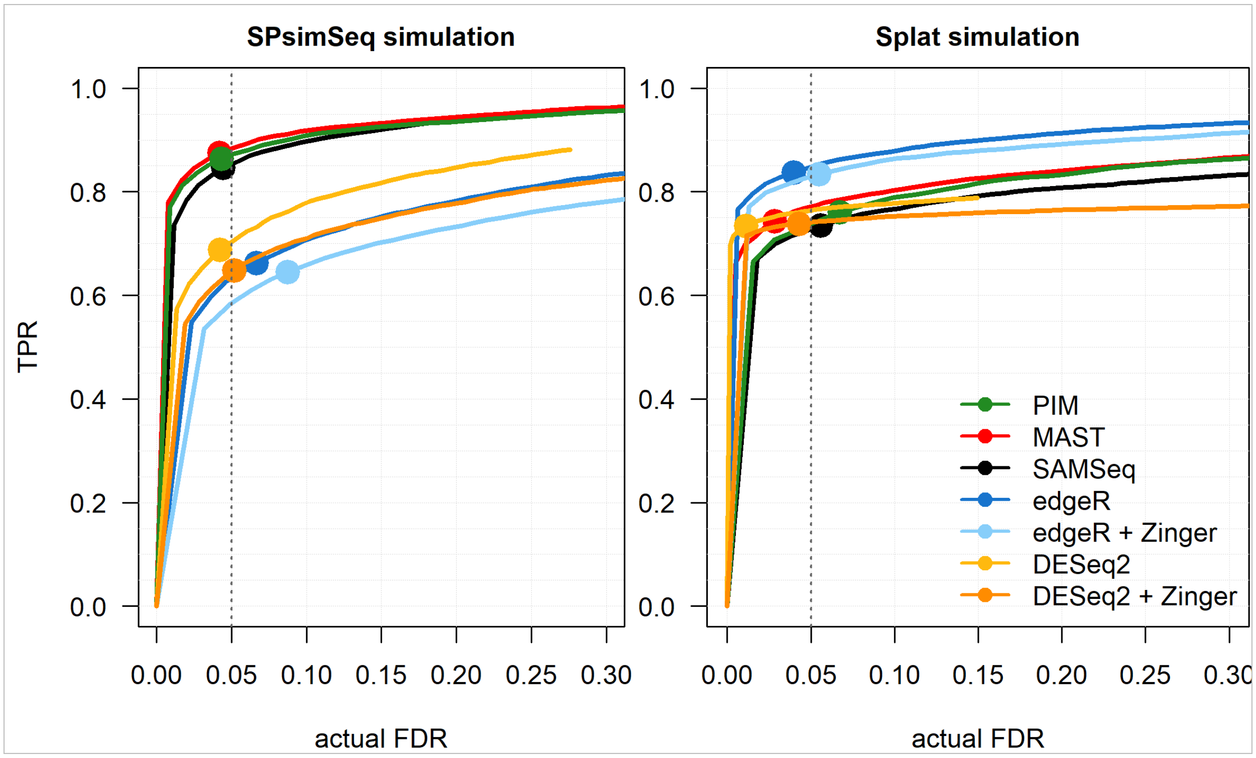

Figure S3: **Results from the simulation study starting from source dataset A.** Dataset A is neuroblastoma scRNA-seq data generated with SMARTer/C1 protocol. Each simulated dataset includes 2500 genes among which 10% DE, and 2 experimental groups with each 100 cells. For the SPsimSeq simulation, the DE genes have LFC $\geq 1$ in source data (data A), whereas for the Splat simulation the FC for DE genes is sampled from a log-Normal(location=1.5, scale=0.4), such that more than 97.5% of the DE genes have a LFC of at least 1. The performance measures (actual FDR and TPR) are averaged over a total of 50 independent simulation runs. The curves represent actual FDR and TPR evaluated at nominal FDR levels ranging from 0 to 0.3, and the solid dots show the performances at the 5% nominal FDR level.

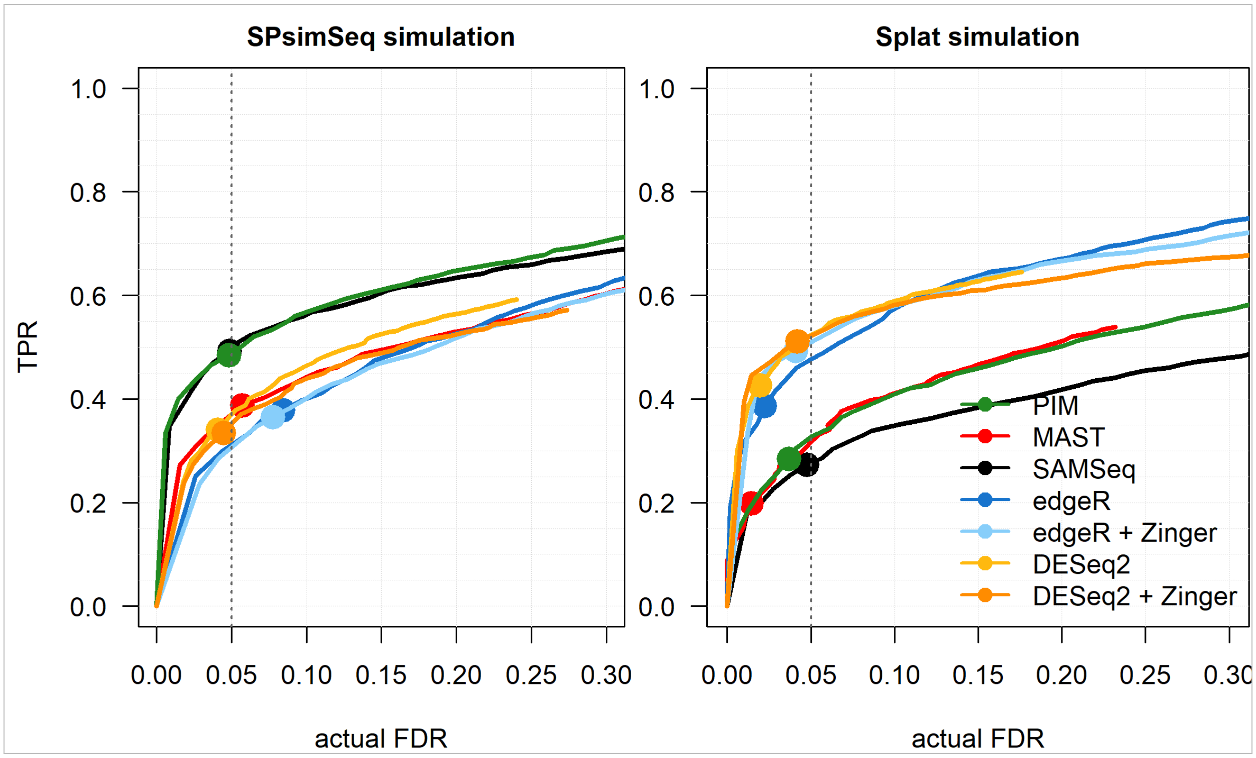

Figure S4: **Results from the simulation study starting from source dataset A.** Dataset A is neuroblastoma scRNA-seq data generated with SMARTer/C1 protocol. Each simulated dataset includes 2500 genes among which 10% DE, and 2 experimental groups with each 100 cells. For the SPsimSeq simulation, the DE genes have LFC $\geq 0.5$ in source data (data A), whereas for the Splat simulation the FC for DE genes is sampled from a log-Normal(location=1.25, scale=0.4), such that more than 97.5% of the DE genes have a LFC of at least 0.5. The performance measures (actual FDR and TPR) are averaged over a total of 50 independent simulation runs. The curves represent actual FDR and TPR evaluated at nominal FDR levels ranging from 0 to 0.3, and the solid dots show the performances at the 5% nominal FDR level.

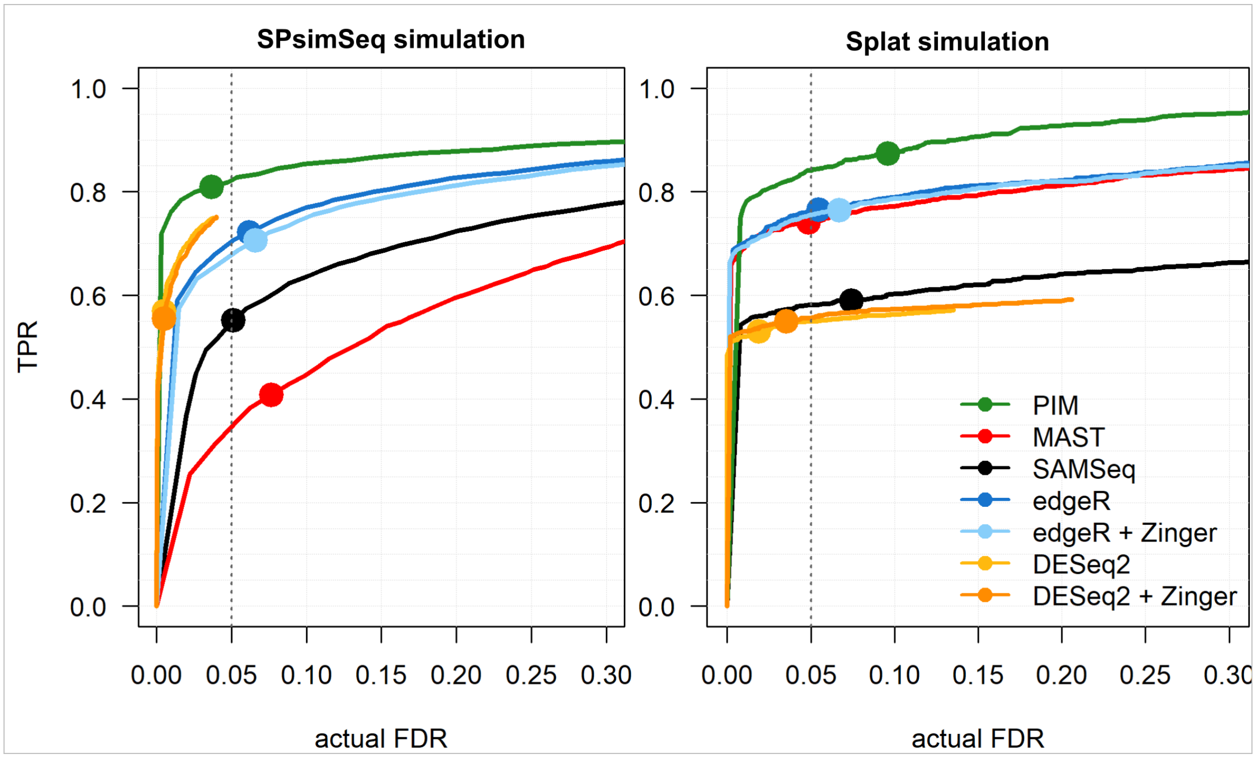

Figure S5: **Results from the simulation study starting from source dataset B.** Dataset B is neuroblastoma scRNA-seq data generated with Chromium protocol (UMI counts). Each simulated dataset includes 2500 genes among which 10% DE, and 2 experimental groups with each 100 cells. For the SPsimSeq simulation, the DE genes have LFC $\geq 1$ in source data (data B), whereas for the Splat simulation the FC for DE genes is sampled from a log-Normal(location=1.5, scale=0.4), such that more than 97.5% of the DE genes have a LFC of at least 1. The performance measures (actual FDR and TPR) are averaged over a total of 50 independent simulation runs. The curves represents actual FDR and TPR evaluated at nominal FDR levels ranging from 0 to 0.3, and the solid dots show the performances at the 5% nominal FDR level.

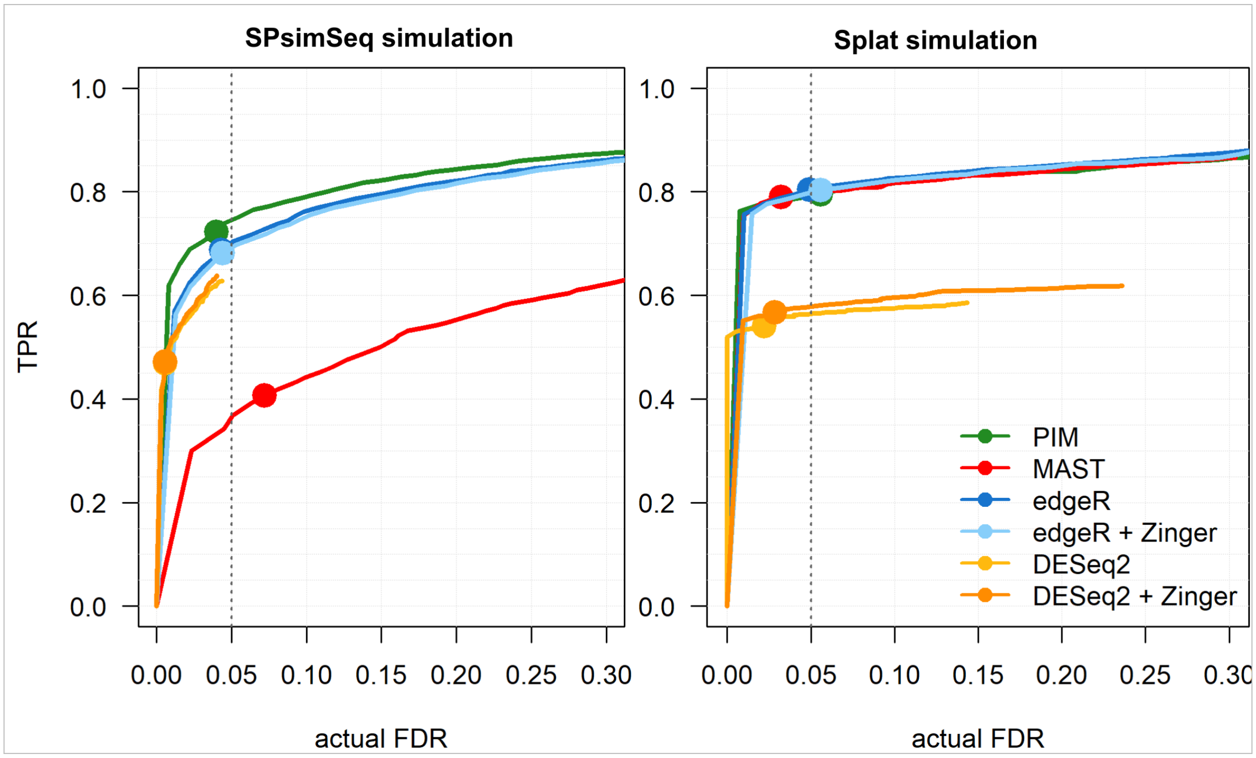

Figure S6: **Results from the simulation study starting from source dataset B.** Dataset B is neuroblastoma scRNA-seq data generated with Chromium protocol (UMI counts). Each simulated dataset includes 2500 genes among which 10% DE, and 2 experimental groups with each 200 cells. For the SPsimSeq simulation, the DE genes have LFC $\geq 0.5$ in source data (data B), whereas for the Splat simulation the FC for DE genes is sampled from a log-Normal(location=1.25, scale=0.4), such that more than 97.5% of the DE genes have a LFC of at least 0.5. The performance measures (actual FDR and TPR) are averaged over a total of 50 independent simulation runs. The curves represents actual FDR and TPR evaluated at nominal FDR levels ranging from 0 to 0.3, and the solid dots show the performances at the 5% nominal FDR level. SAMSeq failed for datasets simulated in this simulation setting and exluded from the result.

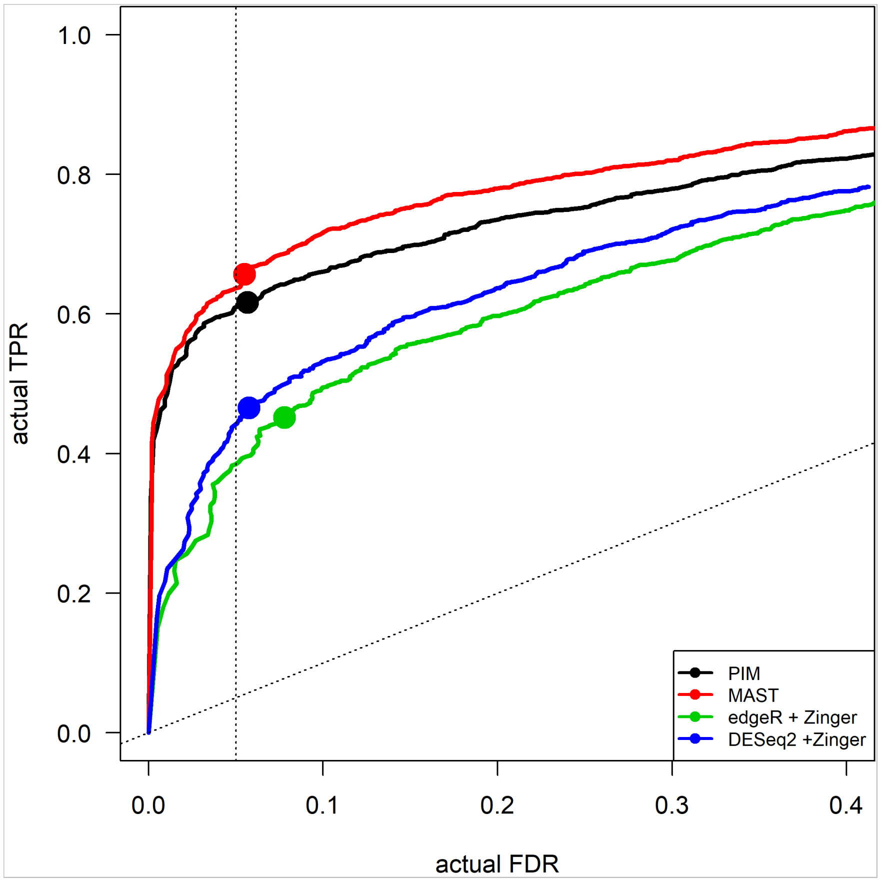

Figure S7: Performance of DE tools for testing DGE across three independent groups of cells based on the SPsimSeq simulation starting from dataset A . In particular, three groups of cells were generated by partitioning the simulated control group (vehicle) into two mock groups. The objective is to demonstrate the applicability of PIM in multiple group designs and evaluate its performance in comparison to the other regression based parametric tools. Each simulated dataset includes 2500 genes, among which 10% DE, and 3 experimental groups, each with 50 cells. The DE genes have LFC $\geq 0.5$ between the control and the treatment group. The actual FDR and TPR are calculated by averaging over a total of 30 independent simulation runs. The curve represents actual FDR and TPR evaluated at different nominal FDR levels, ranging from 0 to 0.5, and the solid dots show the performances at the 5% nominal FDR level.

#### 2.2 Mock comparison

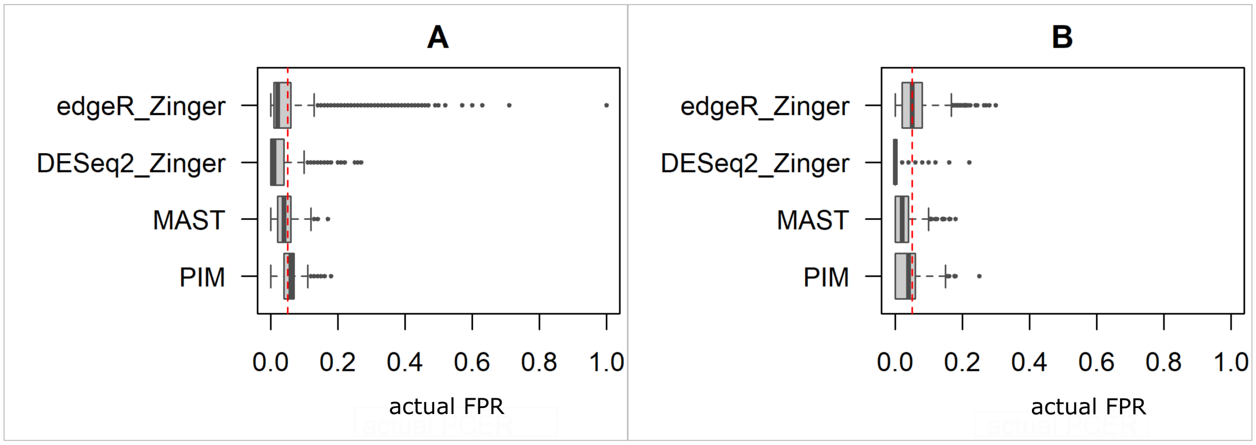

Figure S8: The actual false positive rate (FPR), which is defined for each gene as the fraction of simulations with unadjusted p-value less than 5% (nominal PCER=5% and indicated by red dashed vertical line), as calculated from the mock simulations, for datasets A (panel A) and B (panel B). The box plot shows the distribution across genes.

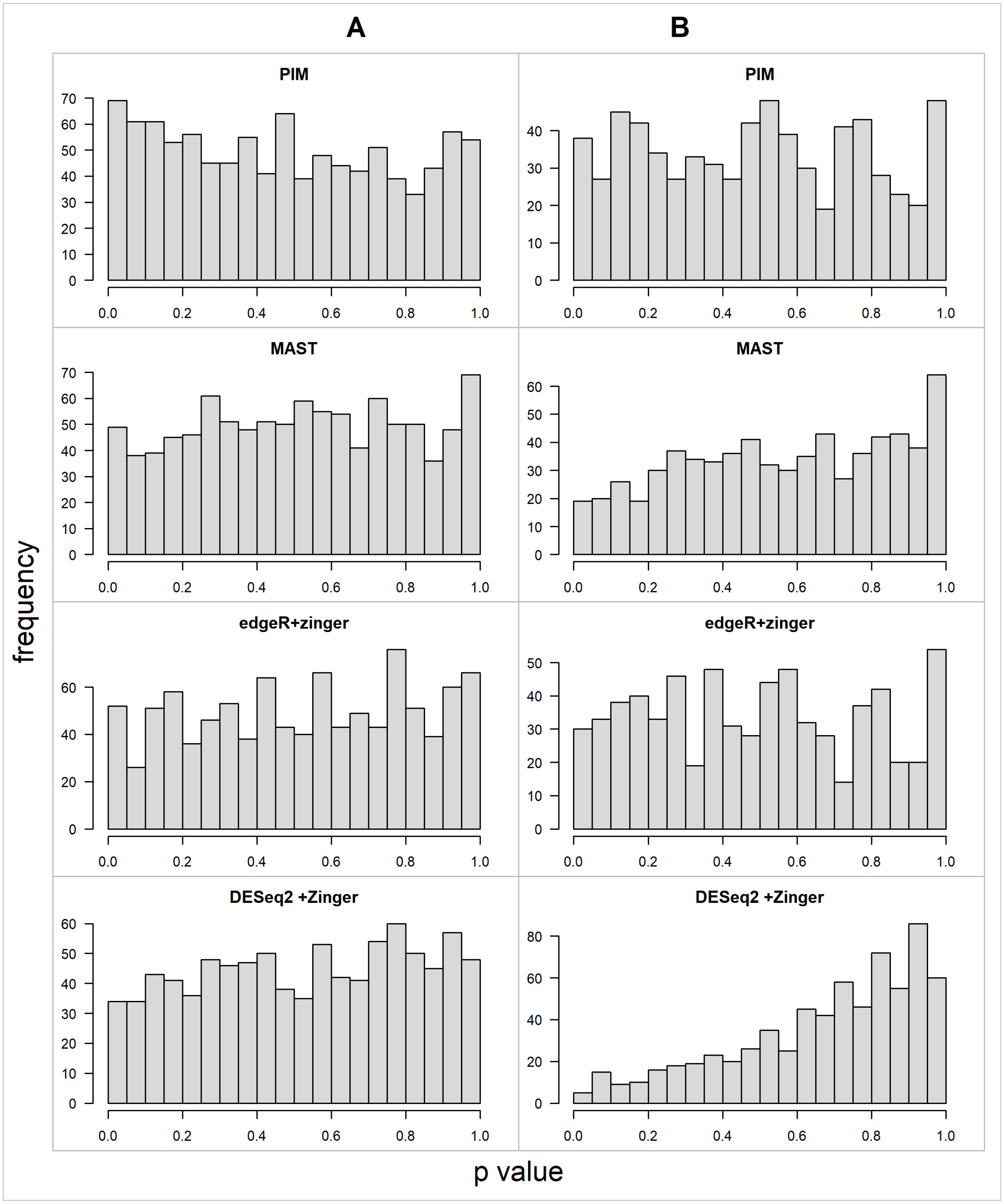

Figure S9: Distribution of unadjusted p-values from mock simulations, starting from source datasets A (left) and B (right).

### 3 Additional results from analysis of real scRNA-seq datasets

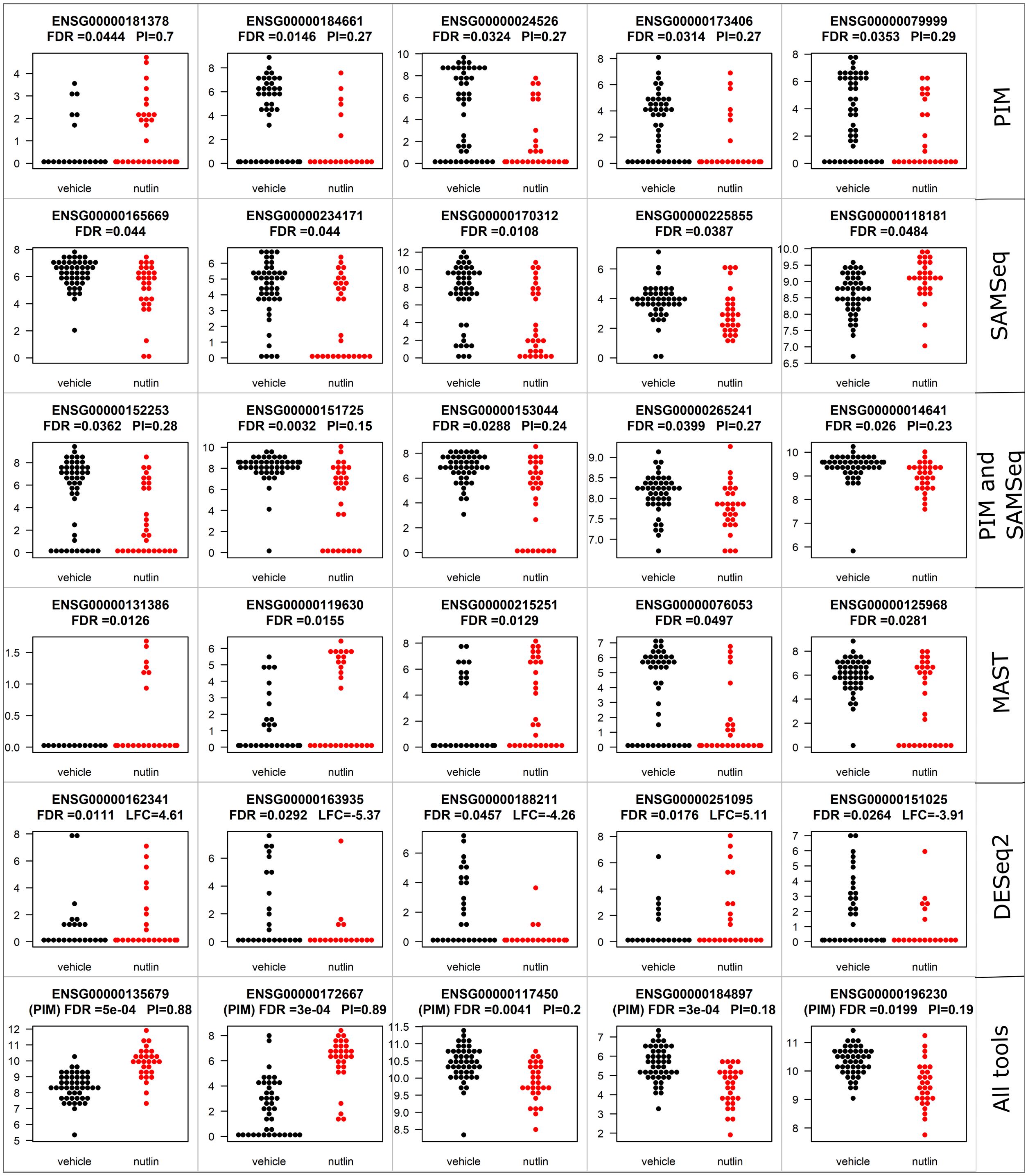

Figure S10: Beeswarm plot to demonstrate the distribution of log-CPM for 5 randomly selected genes among those that are uniquely called DE by PIM, SAMSeq, PIM and SAMSeq, MAST, and DESeq2 at the 5% FDR level and from commonly detected DE genes (bottom panel). The distributions are shown for each group (nutlin and vehicle) separately.

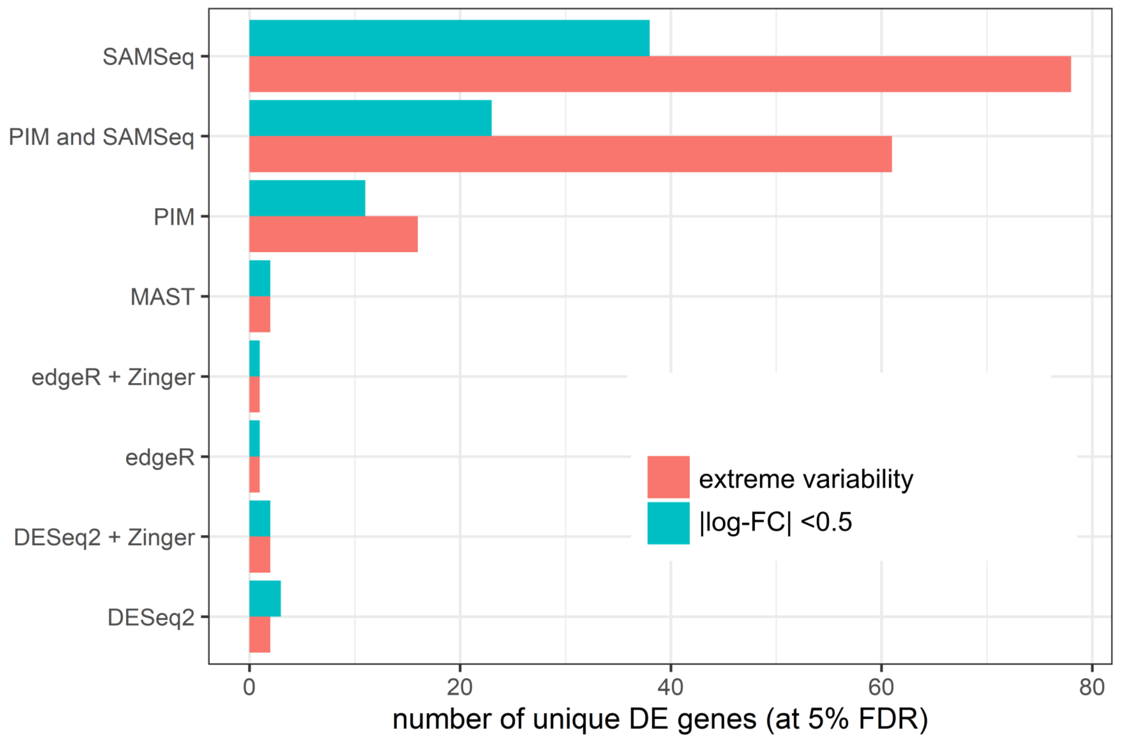

Figure S11: The characteristics of genes uniquely called as DE (at 5% FDR) by each tool. The characteristics include variability CPM across cells and the log fold change estimates. For each tool we counted the number of uniquely identified DE genes that have extreme variability (the upper 10%), and those with estimated log-fold-change below 0.5.

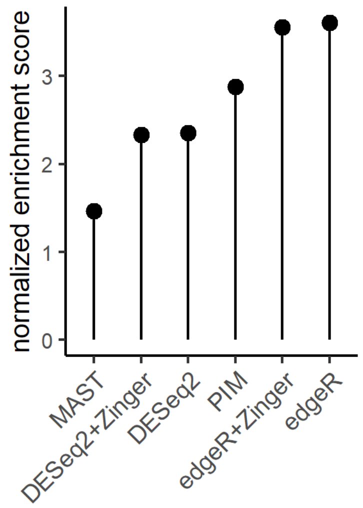

Figure S12: GSEA results for dataset B.
